# Structure of *E. coli* Twin-arginine translocase (Tat) complex with bound cargo

**DOI:** 10.1101/2025.09.16.676506

**Authors:** Ziyu Zhao, Leonid A. Sazanov

## Abstract

How the twin-arginine translocase (Tat) system transports fully folded substrate proteins across cellular membranes without disrupting membrane integrity has been a fundamental question in cell biology for decades. The Tat system recognizes cargo signal peptide via a conserved twin-arginine motif and is found in prokaryotes and plant organelles. Multi-subunit Tat complex facilitates proton motive force-dependent translocation process, yet its overall architecture remains unknown. Here, we present an atomic cryo-EM structure of a *E. coli* trimeric TatB_3_C_3_ complex bound to the substrate SufI. The complex adopts an unusual wide-open, bowl-shaped architecture with a polar inner cavity. Unexpectedly, the cargo is engaged in a dual-contact mode: while the signal peptide binds inside one TatBC unit, the folded domain docks tightly onto an adjacent unit. The structure offers a mechanistic framework for substrate engagement and translocation by the Tat system, suggesting a direct involvement of the entire Tat complex in substrate translocation.

## Introduction

The twin-arginine translocase (Tat) (*1*–*3*) mediates protein transport across membranes in bacteria, archaea and plant organelles. Unlike the Sec system (*4*), which translocates unfolded proteins, Tat uniquely exports fully folded proteins out of cytoplasm (*5*). In bacteria, Tat translocase is essential for metabolic processes (*5*) and virulence (*6*), and so is an important antimicrobial drug target. In plants it is essential for photosynthesis (*7*).

In *E. coli*, Tat complexes consist of multiple copies of TatA, TatB and TatC subunits. TatC is a glove-shaped protein comprising six transmembrane helices (TMH) (*8, 9*). TatA and TatB share a similar structure, each consisting of a short N-terminal TMH followed by an amphipathic helix (APH), which is typically longer in TatB than in TatA (*10, 11*). Cargo proteins are targeted to Tat via N-terminal signal peptides with a conserved twin-arginine motif. Initial binding at the cytoplasmic surface triggers the recruitment of additional TatA subunits forming an activated translocase that moves cargos into periplasm. This process is driven by the proton motive force (*pmf*) (*12, 3*).

How the Tat system achieves a feat of moving fully folded bulky proteins across membranes while maintaining membrane integrity is one of mysteries of biology. Although atomic structures of individual TatA, TatB and TatC subunits have been defined (*8*–*11*), the architecture of the assembled Tat complex remains unknown despite intensive efforts. Here we present a cryo-EM structure of the *Escherichia coli* (*E. coli*) TatBC complex in its resting trimeric state bound to the substrate SufI at the overall resolution of ∼3.74 Å. The core of the complex is resolved to about 3.5 Å, allowing the refinement of atomic model. TatB_3_C_3_ complex adopts a cytoplasm-facing, bowl-like architecture, with a charged inner cavity. TatC-TatC interactions at the periplasmic side form the scaffold of the complex, while TatB subunits act as molecular “glue”, stabilizing cytoplasmic side and core. Cargo protein SufI targets to the cytoplasmic side of one TatBC unit and docks onto an adjacent TatBC unit. Our structure provides critical insights into the architecture of TatBC receptor complex and its substrate recognition and translocation mechanism. We propose that cargo proteins are translocated through TatABC complex, rather than through separate TatA oligomers, via expansion of inner cavity.

## Protein expression/purification and data collection/processing

TatABC and SufI were co-expressed in a *tatABC* deletion Top10 *E. coli* strain. For purification of Tat complex, a 3XFLAG tag was fused to the C-terminus of TatC. Functional assays indicated that the addition of tag did not impair Tat translocation activity (fig. S1). In addition, to accumulate SufI in the cytoplasm, a SufI–8×His construct was expressed and immunoblot analysis using anti-His antibody confirmed successful translocation of SufI–His. Elevated Tat expression levels correlated with increased translocation efficiency (fig. S2).

Cells expressing TatABC and saturating levels of SufI were solubilized in a mild detergent digitonin and the Tat complex, containing all three Tat subunits, was initially purified using anti-FLAG resin and size-exclusion chromatography (fig. S3). Tat-containing fractions were subsequently further purified via nickel affinity pull-down to enrich for Tat–SufI complexes (fig. S3). Cryo-EM data were collected using a Titan Krios microscope and a total of ∼10k micrographs yielded ∼1.3 M good particles after 2D classification. 3D classification resulted in a subset of 164 k high-quality particles. Final refinement involved particle subtraction to remove micelle background contribution, yielding final density map. The processing workflow is shown in fig. S4, representative density fitting of subunits in fig. S5, data processing statistics in Table S1 and the summary of modelling in Table S2.

During 2D classification, particles containing either one (∼73% of particles) or two SufI (∼27%) molecules attached to Tat complex were observed (fig. S6), consistent with prior low-resolution maps of TatBC–SufI (*13*). However, only one-SufI containing particles can be aligned properly to produce a high-resolution map. The second copy of SufI is weakly attached, and below we discuss one SufI-bound structure, unless stated otherwise.

## Overall structure of TatB_3_C_3_-SufI complex

The Tat receptor complex is a TatBC trimer, comprising three copies each of TatB and TatC, and one bound SufI molecule (Fig. 1). The *E. coli* TatC comprises six transmembrane (TM) helices, including four long TM helices followed by two shorter TM5-6 (Figs. 1b and 3a). Three TatC subunits are highly tilted relative to membrane, forming a strikingly wide cytoplasm-facing bowl-like architecture, with side lengths of ∼76 Å (Fig. 1). TM1-4 form outer walls of the bowl, while TM5-6 and TatB TMH form an inward-tilted three-helix bundle, nearly parallel to the membrane plane, like amphipathic helices. The APH of TatB proceeds from its TMH to approach TatC TM1 at cytoplasmic side.

**Fig. 1.**
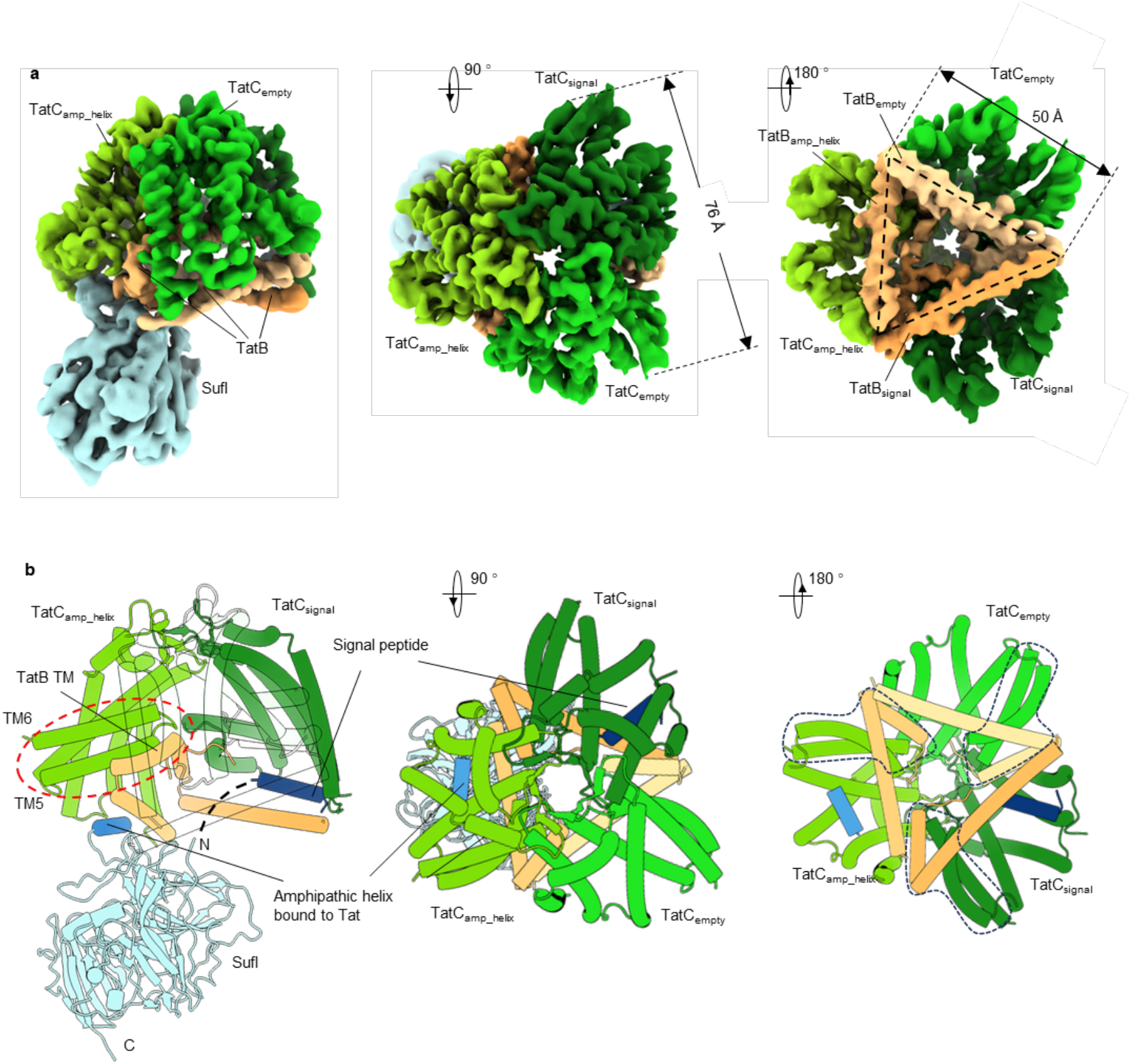
Overall structure of TatB_3_C_3_-SufI complex. **a**, Cryo-EM density map of the TatB_3_C_3_-SufI complex. SufI density is omitted in the right panel for clarity. **b**, Cartoon model of the TatB_3_C_3_-SufI complex. TatC_empty_ and TatB_empty_ are hidden for clarity. TM5 and TM6 of TatC form a three-helix bundle with the short TMH of TatB, highlighted by a red dashed circle. The signal peptide of SufI is shown in dark blue, and the short amphipathic helix SA of SufI interacting with Tat components is highlighted in blue. The middle and right panel show top (periplasmic) and bottom (cytoplasmic) views of the complex. Throughout, TatC subunits are coloured in shades of green, TatB subunits in yellow and SufI in cyan.

SufI engages two distinct TatBC units at the cytoplasmic side (Fig. 1b): its signal peptide binds near the TatC TM1–2 interface of one unit (TatBC_signal_), running beneath the APH of TatB; and a short amphipathic helix (SA helix, SufI residues 301–310) docks onto a neighboring unit (TatBC_amp_helix_), in a roughly similar location between TatC TM1–3 and the APH of TatB. The third TatBC unit (TatBC_empty_) remains unoccupied.

The best-resolved regions lie within the periplasmic and TM domains, while cytoplasmic-facing surfaces appear more flexible (fig. S4). Several regions were not resolved in the cryo-EM map, including the C-terminal helical stretch of TatC, cytoplasmic helices 3-4 of TatB, and SufI residues 18–29, likely due to their high flexibility. The C-termini of TatC and TatB are less conserved (figs. S7 and S8) and may be less important functionally. Although TatA was present during initial purification, it appeared sub-stoichiometric after SufI affinity enrichment (fig. S3b), indicating that some TatA subunits may dissociate from activated Tat complex during purification. No TatA density was observed in the cryo-EM map, suggesting it was either too dynamic or too sub-stoichiometric to be resolved.

Each TatB N-terminal TMH interacts with and runs anti-parallel to TatC TM5, lying nearly parallel to the membrane (Fig. 1b, left panel). These three-helix bundles (TatB TMH plus TatC TM5-6) meet in the centre, stabilizing the open bowl-like architecture (Fig. 1b, right panel). The three N-terminal loops of TatB form an orifice in the cavity center, located under the blocked TatC periplasmic pore (Figs. 1 and 4b). The amphipathic helices of TatB sit on the cytoplasmic surface of the complex and create an inner triangle with side lengths of ∼50 Å, partly covering the inner cavity and stabilizing the structure (Fig. 1, right panels).

TatBC complex has a sealed periplasmic face, as would be expected to prevent leaks across the membrane (Fig. 2bc). However, the central seal, where the three TatC subunits meet, is very thin. The potential pore here is blocked mainly by just one residue – a conserved TatC V60, along with T62 (figs. S7 and S9). This is energetically unfavorable for a permanent membrane sealing, suggesting that this central pore may be able to open during the translocation cycle. Mutations of I60 and nearby residues abolish transportation activity (*14*) (Table S4), highlighting the importance of these conserved residues. The observed loss of transport activity is likely due to disruption of the local environment required for conformational changes in Tat components and/or break of the fragile central seal.

**Fig. 2.**
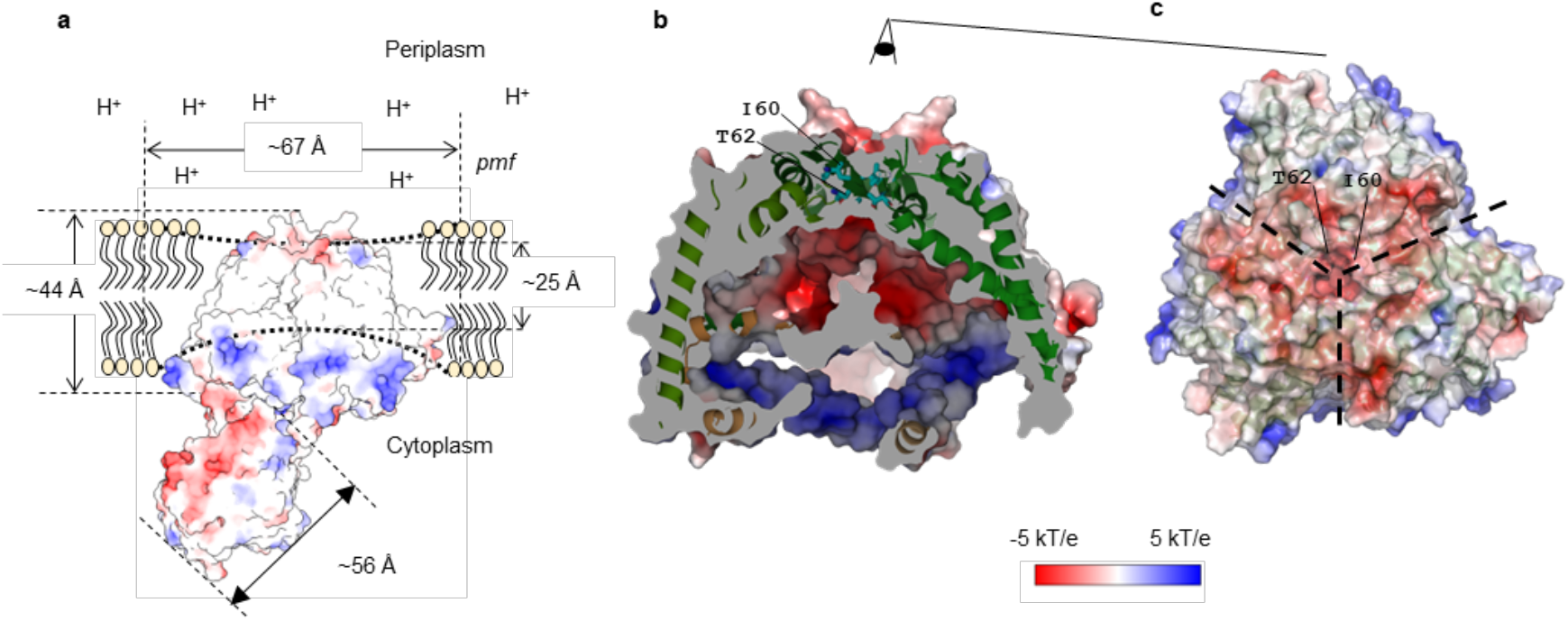
Electrostatic surface potential and membrane integration of the TatB_3_C_3_ complex. **a**, Solvent-excluded surface, colored according to electrostatic potential, showing the orientation of the complex within the membrane and highlighting its contribution to a locally thinned membrane region. The overall height of the complex is approximately 44 Å, with the hydrophobic transmembrane region spanning ∼25 Å. *pmf* represents proton motive force. **b**, Cross-sectional view showing the solvent-accessible surface, colored according to electrostatic potential, within the inner cavity of the complex. TatC I60 and T62 sealing the periplasmic side of Tat are shown as sticks. **c**, Top view of the complex showing the periplasmic surface. TatC I60 and T62 contribute to sealing the periplasmic side of the cavity.

The total height of TatBC (∼44 Å) is consistent with average membrane thickness (Fig. 2a). However, the exposed hydrophobic surface is only ∼25 Å thick, due to tilted TatC TM helices, indicating local membrane thinning, potentially lowering energy barrier for substrate translocation. The periplasmic surface of TatBC is negatively charged, as is common for bacterial membrane proteins (Fig. 2c). The cytoplasmic side is positively charged (Fig. 2b), attracting the negatively charged face of SufI (Fig. 2a). The inner cavity is negatively charged (Fig. 2b), indicating that it is not filled with lipids and may be available for substrate pass through, if it is accompanied by appropriate conformational changes. TatC-D63, D150, E170 and D211 contribute to the negative charge of inner cavity (fig. S9). Mutation of any of these residues lead to abolished or reduced transportation (*15, 16, 14*), highlighting the importance of charged cavity for transportation (Table S4).

Based on these findings, we propose that our structure reflects a resting state of the Tat complex, in which SufI is poised for translocation but has not yet engaged recruitment of additional TatA subunits.

## *E. coli* TatB and TatC structures

TatC_signal_, TatC_helix_ and TatC_empty_ exhibit overall similar structures (Fig. 3a), with slight deviations at the C-terminus of TM4. While the overall architecture of *E. coli* TatC subunit resembles that of crystal structure of *Aquifex aeolicus* (*Aquifex*) TatC monomer (*8*) (Fig. 3b), notable differences are present. In particular, TM3 of *E. coli* TatC is oriented more vertically relative to the membrane plane, whereas TM5–6 adopt much more tilted configuration (Fig. 3bd), likely due to interaction with TatB TMH. On the periplasmic side, *E. coli* TatC forms parallel β-sheet interactions between residues M59–A61 and T149–I151 of one subunit, as well as between M59–A61 and V145– V147 of an adjacent TatC subunit, thus forming a short parallel β-barrel in the centre (Figs. 3c and 1b). The apparent rigidity of this architecture may help to keep the central pore blocked in the resting state (Fig. 2bc). However, the parallel ß-sheet interactions are less stable than anti-parallel, and this design possibly facilitates the later disruption of these interactions to allow cargo passage. The corresponding region in *Aquifex* monomer differs in structure (Fig. 3c) (*8*), but may adopt a similar conformation upon complex formation.

**Fig. 3.**
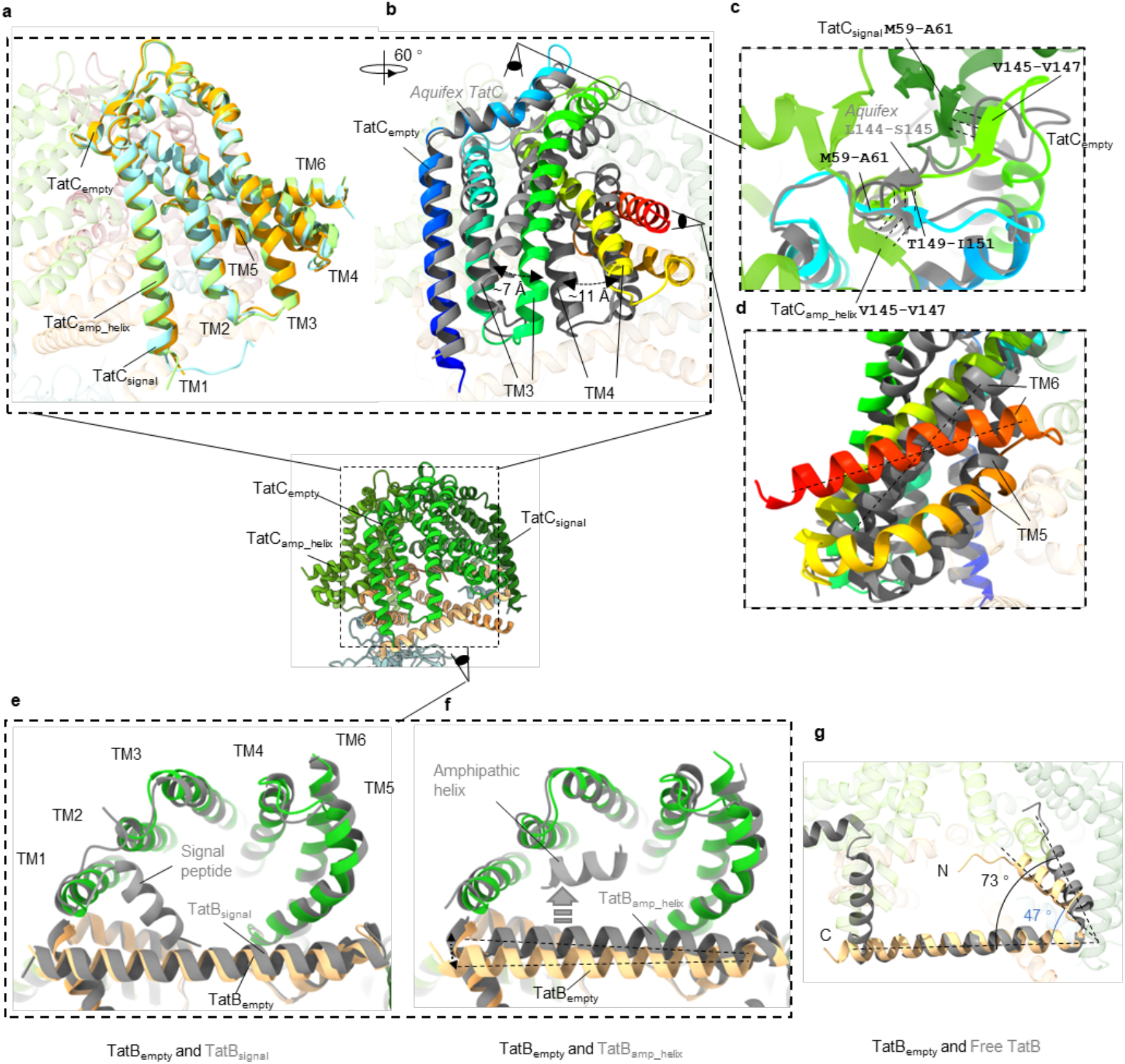
Structural features of *E. coli* TatC and TatB. Comparisons of *E. coli* TatC subunits with each other, with *Aquifex* TatC crystal structure (PDB ID: 4B4A) and with NMR structure of *E. coli* TatB (PDB ID: 2MI2). The central panel shows the overall structure of the TatB_3_C_3_–SufI complex, highlighting the orientations used to depict specific interaction interfaces. **a**, Structural alignment of TatC_empty_ (orange), TatC_amp_helix_ (light green), TatC_signal_ (cyan). **b**, Superimposition of *E. coli* TatC_empty_ (coloured from blue to red N-to-C-terminus) and *Aquifex* TatC (grey), highlighting structural consensus and differences. **c**, Top view of TatC_empty_ overlaid with *Aquifex* TatC (grey). Parallel β-sheet hydrogen bonds at the periplasmic interface of TatC are indicated with dashed black lines. **d**, Side view comparison of TM5 and TM6 between *E. coli* TatC_empty_ (coloured from blue to red N-to-C-terminus) and *Aquifex* TatC (grey), showing distinct helix tilting within TatBC complex. **e, f**, Bottom views of TatB_empty_ superimposed with TatB_signal_ and TatB_amp_helix_. **g**, NMR TatB (PDB ID: 2MI2) structure (grey) is superimposed with TatB_empty_, showing angular difference of short TMH and amphipathic helix.

**Fig. 4.**
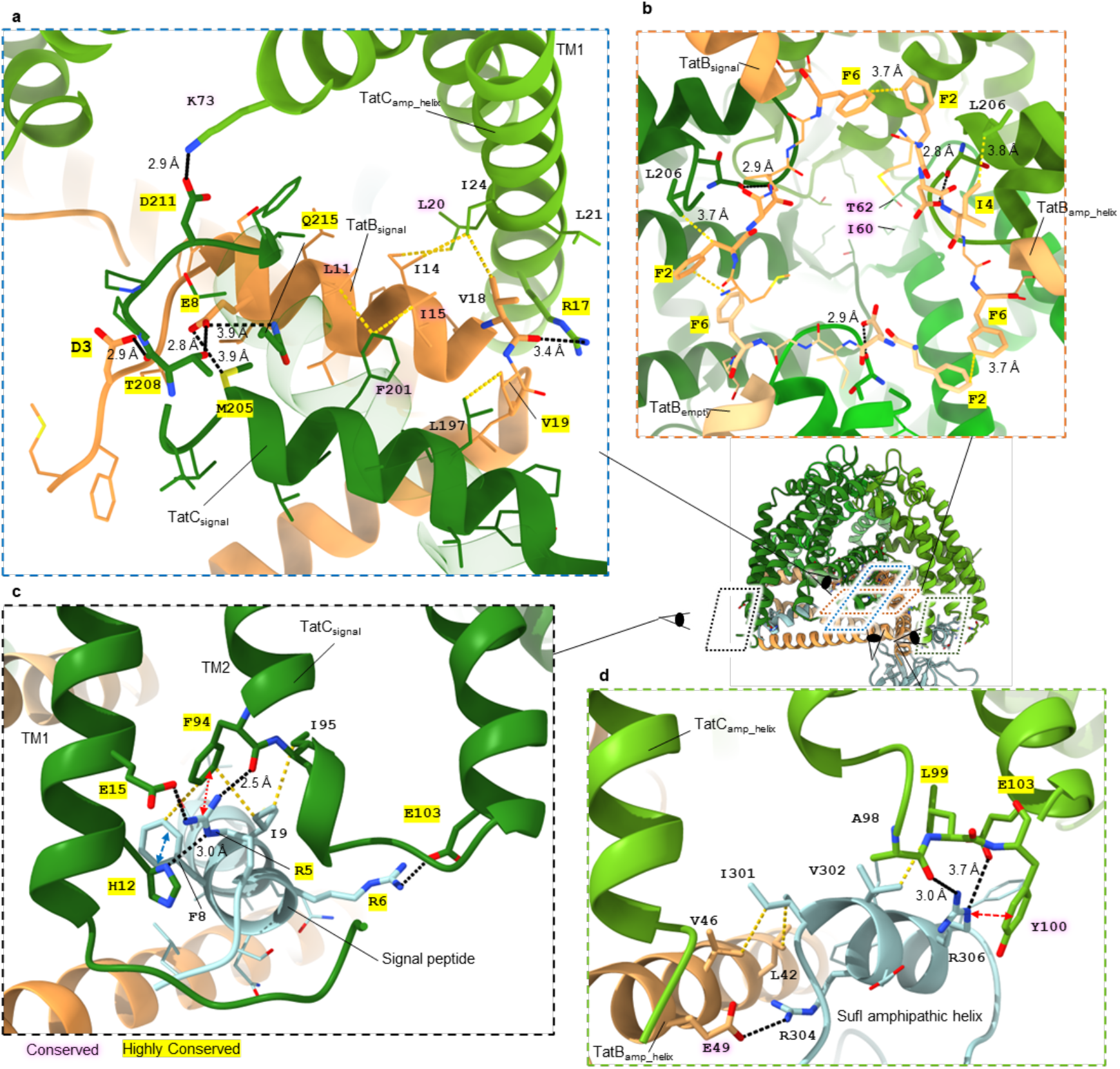
Interactions within TatBC and between SufI and Tat complex. The central right panel shows the overall structure of the TatB_3_C_3_–SufI complex, highlighting the orientations used to depict specific interaction interfaces. H-bonds/salt bridges are shown as black dash lines, with distances of interactions indicated (except when side-chain density is unclear). Hydrophobic interactions are shown as yellow dashed lines and are between 3.6 Å to 4 Å. Highly conserved residues (ConSurf score 9) are highlighted in yellow, while conserved residues (score 8) are shaded in pink. **a**, Interactions between TatB and TatC. **b**, Bottom view of interactions involving the N-terminal ends of TatB. **c**, Interactions between signal peptide and TatC. A cation–π interaction between SufI R5 and TatC F94 (3.3 Å) is indicated by a red dashed line with arrowheads. A π–π stacking interaction (5.1 Å) is shown by a blue dashed line with arrowheads. **d**, Interactions between the short amphipathic helix of SufI and TatB/TatC. A cation–π interaction between TatC Y100 and SufI R306 (3.5 Å) is indicated by a red dashed line with arrowheads.

TatB forms an asymmetric trimer within the complex. Upon binding of SA helix from SufI, the C-terminal end of APH of TatB_amp_helix_ shifts closer to the TM2–3 region of TatC, likely due to interactions with SufI (Fig. 3f). On the other hand, binding of signal peptide does not induce major changes in interacting TatB and TatC regions (Fig. 3e). Compared to its NMR structure in the unbound state (*11*), TatB adopts a more compact conformation in the complex, with the angle between TMH and APH reduced from ∼73° to ∼47° (Fig. 3g). This suggests that TatB is flexible and may undergo rearrangement during substrate translocation.

Interestingly, TatB F13 adopts different conformations in TatB_signal_ and TatB_amp_helix_ states (fig. S10). In TatB_signal_, it stabilizes the cytoplasmic side via hydrophobic contacts with TatC L16 and L20 (fig. S10), while in TatB_amp_helix_, these interactions are lost. The F13Y mutant, known to activate Tat without signal peptide (*17*) (Table S4), likely weakens these interactions and possibly lowers the activation barrier.

## Scaffolding TatC-TatC interactions

TatC subunits primarily interact with each other on the periplasmic side of the complex through parallel β-sheet hydrogen bonding noted above (Figs. 3c and 1b). In addition, TatC monomers engage in hydrophobic interactions at their TM interfaces on the periplasmic side (fig. S11a). Substitution of these hydrophobic residues with polar amino acids (V64E, F68S and L137H) disrupts Tat-dependent transport (*14*) (Table S4), suggesting that these interactions contribute to the structural stability of the Tat complex.

A salt bridge is observed between the conserved D211 and K73 from the adjacent TatC subunits (Figs. 4a and S11b). Mutation of TatC D211 to alanine or deletion of K73 abolish transport activity (*14, 15*) (Table S4), indicating that this salt bridge is critical for stability of TatBC complex. These bridges likely lock the central three-helix bundles (TatC TM5,6 and TatB TMH) in the position to form the central iris-like orifice (Fig. 1b). The charge state of these bridges may depend on *pmf*, potentially contributing to any *pmf*-driven conformational changes. Additionally, this key K73 sits on TatC TM2, which is broken and bent in the middle by the conserved P85 (fig. S11b). P85A mutation significantly reduces translocation efficiency (Table S4) (*15*). This mutation possibly prevents periplasmic sealing of core complex and coordination between TatC TM2 and central three-helix bundles.

In contrast, no interactions are observed between TatC monomers on the cytoplasmic side, which is instead stabilised by TatB subunits (Fig. 1). This absence of inter-TatC contacts suggests a more flexible and dynamic cytoplasmic region, in agreement with location resolution map (fig. S4), potentially facilitating TatA and TatB exchange (*18*) or enabling TatA recruitment during activation and translocation.

## TatB-TatC interactions “glue” TatC subunits together

The short TMH of TatB closely associates with TM5 of TatC via hydrophobic interactions (Fig. 4a and 1b), consistent with previous analyses (*14, 19, 20*). Additionally, the TatB E8 H-bonds to TatC M205, T208, and Q215 (Fig. 4a). Interactions with these conserved polar cluster (MTQ) residues promote the binding of TatB to TatC and successful cargo transportation (*20*) (Table S4). In the hydrophobic environment of the membrane, these polar interactions likely serve to pin TatB in the correct position relative to TatC TM5, preventing lateral movement and contributing to the structural integrity of the Tat complex.

Moreover, the conserved TatB D3 may form H-bond with TatC T208 backbone oxygen, provided that aspartate is protonated, which is more likely in a hydrophobic environment (*21*). Furthermore, the side chains of TatC L206 and TatB F2, I4 are likely to form hydrophobic interactions (Fig. 4b). Overall, the TatB N-terminal loop and TatC TM5-6 loop appear to interact strongly, stabilizing the central orifice area.

TatB also interacts with adjacent TatC via hydrophobic and polar interactions (Fig. 4a). Specifically, I14 and V18 from TatB TMH form hydrophobic contacts with L20 and I24 on TM1 of the neighboring TatC. Conserved TatC R17 from TM1 of the neighboring TatC neutralizes negative dipole charge of the exposed C-terminus of TatB TMH. Mutations of TatC R17 impair transport activity and destabilize the Tat complex (*15, 22, 23*) (Table S4), underscoring its critical role in trimerization of the resting state of Tat complex.

In addition, the C-terminal end of TatB APH likely interacts with the N-terminus of TatC TM1 and with the N-terminal part of neighboring TatB APH (fig. S12). Thus, overall, TatB subunits act as molecular “glue”, joining the cytoplasmic sides of TatC subunits.

Further analysis of TatB (and potentially TatA) role in the complex can be facilitated using AlphaFold3 (AF3) predictions. AF3 was able to model mostly correctly the arrangement of TatB_3_C_3_ complex (fig. S13), allowing for some degree of confidence in other AF3 models of different TatABC-SufI assemblies that we produced. Interestingly, in TatC_3_ or TatA_3_C_3_ models TatC TM5-6 were aligned to TM1-4 (as in *Aquifex* crystal structure) (figs. S14 and S15), and only tilted parallel to membrane in TatB-containing models (figs. S13 and S21), likely due to close interaction with TatB TMH. The presence of TatB (and associated interacting three-helix bundles) appears to tighten the bowl, which is even wider in its absence (in TatC_3_ and TatA_3_C_3_ models) (figs. S13-15). TatB TMH also partially fills the interfaces between TatC subunits at the membrane-exposed surface. However, AF3 model of one SufI bound to TatB_3_C_3_ (or TatA_3_B_3_C_3_) did not orient the substrate correctly. The model of three SufI bound to TatA_3_B_3_C_3_ placed SufI in the broadly correct orientation (fig. S13), but the signal peptide and helix SA were predicted to interact with the same TatBC unit, in disagreement with our data.

## TatB-TatB interactions stabilize complex core

TatB subunits interact with one another via their N-terminal tails through hydrophobic interactions (Fig. 4b). Side chains of conserved F2 and F6 from neighboring TatB subunits pack closely together, forming a triangular hydrophobic core of the central orifice of the TatBC trimer. These inter-TatB interactions were observed in previous cross-linking studies (*19*) (Table S4).

TatA possesses conserved N-terminal residues that are less hydrophobic than those in TatB (figs. S8 and S16), suggesting that if TatA replaces TatB during substrate translocation (*18*), these hydrophobic interactions may weaken, thereby facilitating structural rearrangement and inner cavity expansion. AF3 predictions using only TatA and TatC (and SufI) yield a more loosely packed complex (fig. S15) and with incorrect (unbound) SufI positions. Predicted binding of TatA TMH to TatC TM5 is similar to TatB mode, consistent with co-evolutionary analysis (*20*). However, TatA is perhaps less able to tilt TatC TM5-6 towards the center of the trimer like TatB does (fig. S15), preventing the formation of the central orifice. APHs of TatA are too short to interconnect N-terminus of TatC subunits and instead point towards the cytoplasm (fig. S15, S17). Thus, the replacement of TatB with TatA could “open up” the complex, possibly reflecting a state ready to accept a substrate (to be translocated) inside the widened central cavity of the complex.

The N-terminal tail interactions of TatB, combined with interactions with polar cluster, likely induce a bent conformation of TatB TMH compared to its unbound state (Fig. 3g). The N-terminus of TatB from F2 to L9 is highly conserved (fig. S8). It plays a key role in TatB-TatC and TatB-TatB interactions and is likely essential for stability of the resting state of Tat complex. Overall, the orifice structure formed by the centrally joined three-helix bundles likely stabilizes the unusually wide-open bowl-like shape of the complex. It may also represent a “sensor” for the incoming substrate, triggering conformational changes required for translocation.

## Signal peptide recognition and binding

The signal peptide sequence of Tat substrates features a positively charged n-region with twin-arginine motif, a hydrophobic h-region and a c-region with a characteristic AXA cleavage site (*5*). The SufI signal peptide n-region helix (residues 3-16) binds inside the TatC cavity between TM1 and TM2, nearly parallel to the membrane (Fig. 1b and 4c), rather than on the outer surface (*8*) or inserted deeply in a “hairpin” configuration (*24*) as previously proposed. To reach this site, the N-terminus of the substrate passes under the APH of TatB (Fig. 1b and 4c). This internal binding possibly prepares the substrate for entry into the central cavity. Subsequent replacement of TatB with TatA (*18*) can further open up the entrance (fig. S15).

SufI R5 from the twin-arginine motif interacts with several conserved TatC residues (Fig. 4c). It forms H-bonds with H12 and with previously identified signal peptide recognition residue E15 (*8*). R5 side chain also forms an H-bond with the backbone oxygen of F94 and engages in a cation–π interaction with benzyl group of this residue. Mutations of H12 and E15 to alanine reduce transportation efficiency (*15*). Mutations of F94 to alanine and leucine abolish transport activity, while the F94Y mutation retains functionality (*15*) (Table S4), emphasizing the importance of this cation–π interaction in signal peptide binding.

SufI R6 from the twin-arginine motif was not fully resolved in the structure but is predicted by AF3 to form a salt bridge with TatC E103, which was suggested as signal binding site (*8*) (Fig. 4c). The conserved phenylalanine residue following the twin-arginine from the SRRx**F**LK motif, F8, engages in hydrophobic interactions with TatC F94 and in π–π stacking with TatC H12 (Fig. 4c). Additionally, the side chain of SufI I9 lies close to TatC F94 and I95, forming further hydrophobic contacts. While F8L mutation of signal peptide does not impair Tat function, substitutions F8Y, F8A and I9A significantly reduce translocation efficiency (*25*), underlining the importance of these hydrophobic interactions between the signal peptide and TatC on the cytoplasmic side.

The signal peptide residues A17 to A27 (h- and c-regions) and subsequent residues A28, G29 are not observed in our structure, suggesting that this region is flexible (Figs. 1b). High flexibility at this area would be necessary to allow movement of h- and c-regions while the n-region remains “clamped” by TatB APH and TatC TM1-2 during translocation (*26*). The missing 13 amino acids are sufficient for connection from the mature domain of the substrate to the n-region (fig. S18). The h-region and c-region residues could be positioned with SufI V21 near TatC TM5 and subsequent residues adjacent to TatB TMH, consistent with results of Hamsanathan and colleagues (*24*). However, the n-region helix of the SufI signal peptide is oriented more parallel to the membrane bilayer (Figs. 1b and 4c), rather than forming the predicted “hairpin” conformation (*24, 27*–*29*). This indicates that the n-region of signal peptide does not insert as deeply into the membrane at the resting state as previously thought (*24, 28*). Alternatively, n-region of signal peptides from different substrates may engage the Tat complex in distinct ways, although it seems unlikely given the conserved SRRxFLK binding motif (*3, 25*).

## Docking of cargo

Beyond the signal peptide, a short amphipathic helix SA of SufI binds to another TatBC unit between the cytoplasmic ends of TatC TM1-3 and the APH of TatB, i.e. in a location similar to signal peptide but at different TatBC unit (Fig. 4d). Helix SA contains two arginine residues, R304 and R306, that mediate interactions with both TatC and TatB. R304 is oriented toward the C-terminal region of TatB APH and may form a salt bridge with conserved TatB E49. The side chain of R306 points toward the cytoplasmic surface of TatC and likely forms H-bonds with E103 and the backbone oxygen of A98. Thus, signal peptide binding residue E103 interacts also with SA helix, suggesting its dual role in both signal recognition and substrate docking. In addition, SufI R306 engages in a cation–π interaction with TatC Y100. Mutation of Y100 to alanine significantly reduces translocation efficiency (*15, 16*) (Table S4), indicating its importance in substrate engagement. Hydrophobic interactions are also found between the conserved TatC L99 and SufI V302. TatC L99A mutation results in reduced transportation efficiency (*15*).

Surprisingly, the sequence of this SA helix is highly variable among substrates (fig. S19 and S20), suggesting that not all substrates contain this helix and, consequently, may not form identical interactions with Tat components. However, when we predicted additional Tat/substrate complex structures using AF3, approximately half of the models showed the substrate docking to the APH of TatB (fig. S21, Table S3). This finding raises the possibility that this docking mode may be more common or even universal across substrates, albeit possibly in variable docking manner.

Interestingly, in nearly all predicted models, the fourth helix of TatB appears to lose its secondary structure and wrap around the substrate (fig. S21; Table S3). This feature may reflect a substrate-proofreading function, as previously proposed (*30*).

The identification of the second signal peptide-independent SufI binding site (via SA helix) suggests that full engagement of a substrate involves direct contact between the mature domain of substrate and Tat complex. In vitro, mature domain has been shown to participate in the proofreading for transportation process, while cargo size and shape influence transportation efficiency (*31, 32*).

Apart from SA helix, additional H-bonds have also been identified between side-chains of TatB_amp_helix_ and backbone of SufI G63 and N193 (fig. S22), suggesting multiple interaction areas for docking. Such extended docking may help stabilize the Tat/substrate complex and could play a regulatory role in the translocation process. The dual-contact mode (signal peptide recognition and SA helix docking) provides a more comprehensive understanding of how the substrate is engaged and poised for translocation.

### Second bound SufI

As noted above, our data indicate that the resting Tat complex can accommodate two substrates simultaneously (fig. S6ab). Structural superposition of the Tat_signal_ and Tat_amp_helix_ sites reveals steric clashes between the signal peptide and SA helix of SufI (fig. S23). This indicates that a single TatC subunit cannot accept both a signal peptide and docking of substrate, so the second substrate must bind to the unoccupied TatC_empty_ via its signal peptide but it cannot dock since there is no free site for SA helix to bind.

The 3D refinement of two-SufI particles with one-SufI mask yields better resolution (∼ 5 Å) than using two-SufI mask, due to flexibility of the second SufI (Fig. S6b, S24). This reveals a helix-like density at the signal binding site which was empty in one-SufI complex (fig. S24). Therefore, the Tat complex likely is capable of handling simultaneously a fully engaged state (signal binding and docking) of one substrate and initial recognition (only signal peptide binding) of a second substrate.

These observations support a queueing mechanism for sequential translocation – while one copy of substrate is being translocated, the second copy is ready to be engaged. Such a mechanism, possibly also operating in plants (*33*), may enhance translocation efficiency under high cargo load.

We also observed minor 2D classes showing potentially three densities on the cytoplasmic side of Tat (fig. S6c). 3D reconstruction of these particles was not possible due to low amounts of particles. These features may correspond to three weakly bound SufI molecules, suggesting an alternative state in which all three TatC subunits are bound by substrates via signal peptides, but without tight docking, as there are no free sites for SA helices engagement.

## Translocation mechanism

Here, we report the atomic cryo-EM structure of the *E. coli* trimeric TatBC complex bound to the substrate SufI, capturing a fully engaged state of the resting Tat complex (Fig. 1). The structure reveals the overall architecture of the TatBC complex, detailed atomic interactions within the Tat components, specific contacts between the signal peptide and TatC, and a novel additional docking interface between SufI and TatBC. A recent paper suggested trimeric state of thylakoid Tat complex (*34*), so it is likely that our structure represents general architecture of Tat resting state in bacteria and plants.

Narrow hydrophobic strip on the surface of the complex suggests that it induces local membrane thinning (Fig. 2a), perhaps facilitating conformational flexibility and reducing energy barrier for substrate translocation. Extremely thin layer of just two residues blocks the potential central pore at the periplasmic side (Fig. 2bc). Along with charged inner cavity of the complex, these features may establish a conductive environment for substrates to travel through the central cavity upon suitable conformational changes. Notably, overexpression of substrate indeed induces conformational rearrangements in the TM5-6 region of TatC (*17, 18, 35*) and TatA C-terminal region (*36, 37*).

AF3 model of the TatA_3_B_3_C_3_-SufI_3_ complex predicts that TatB TMH interacts with TM5 of TatC as in our structure, while TatA TMH interacts with TM6 of TatC, filling apparent gaps between TatC subunits (fig. S13). This is consistent with cross-linking and co-evolutionary analyses, which suggested that TatA interacts with TM6 in resting complex but switches to TM5 in active translocase (*18, 20*). We also tried multiple TatA copies as input for AF3 model, but no reasonable structures were obtained.

If during translocation TatA replaces TatB at TatC TM5 (*18*), central hydrophobic interactions could be disrupted and inner cavity expanded. AF3 models (fig. S15) suggest that this replacement could trigger TatC TM5-6 to tilt outward, reverting to its ‘natural’ conformation seen in *Aquifex* TatC (*8*), and opening the complex for cargo passage. Such a transition could depend on *pmf* and presence of cargo. Moreover, as the APH of TatA is shorter than that of TatB (fig. S17), the C-terminal end of the TatA APH is unlikely to interact with the N-terminal region of TatC. This would further destabilize the cytoplasmic interface of the complex, contributing to the dynamic rearrangements required for translocation (*36, 38, 37, 35, 18, 17*).

Thus, based on our structure, we propose a model for substrate translocation by the Tat system in *E. coli* (Fig. 5a). Initially, the signal peptide of the substrate binds to one TatBC unit within the resting TatABC complex. This is followed by docking of the substrate (via SA helix in case of SufI) onto an adjacent TatBC protomer. Subsequently, TatB is displaced by TatA, leading to widening of the cytoplasmic side of the complex, opening of the central iris-like orifice due to tilting of TatC TM5-6 back toward complex periphery (Fig. 5a). Depending on substrate size, a number of additional TatA subunits are recruited as required to enlarge the size of the channel by inserting additional copies of their TMHs in between TatC subunits. Proton motive force (*pmf*) is required to drive this translocation, possibly by disrupting the salt bridges that stabilize the central orifice formed by the three-helix bundles. The queuing mechanism, reflecting the second substrate bound initially only via signal peptide (figs. S6 and S24) is shown in Fig. 5b.

**Fig. 5.**
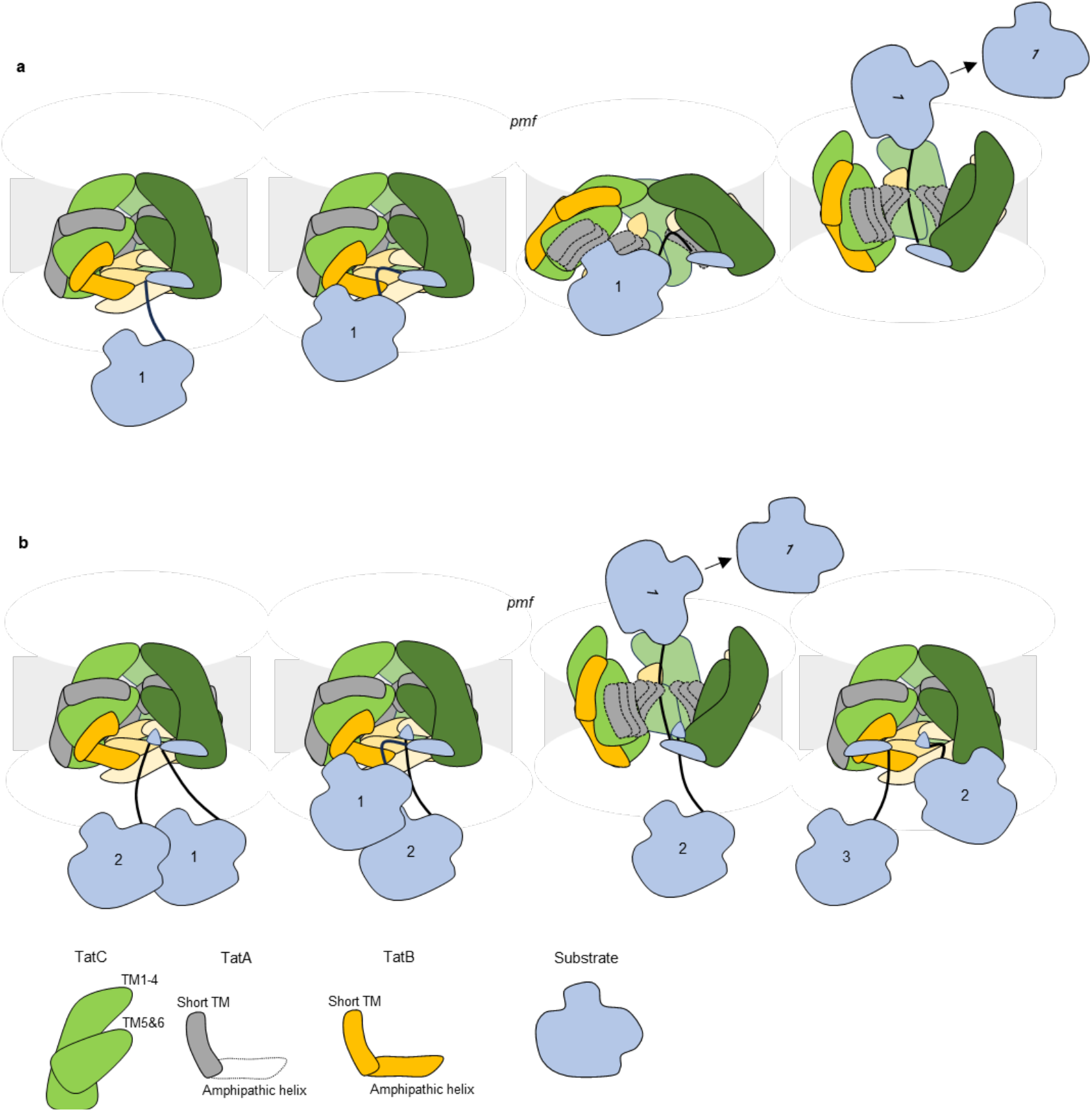
Proposed Tat translocation mechanism. Schematic illustration of the Tat translocation process. Subunit cartoons are shown at the bottom. TatA subunits are depicted in grey to indicate their potential locations according to AF3 models. TatA within the activated translocase is outlined with a dashed line to reflect a putative arrangement, and amphipathic helix of TatA oligomer is not shown. The membrane is shown in light gray. **a**, In the resting state, the signal peptide of the substrate (cargo) initially binds to the cytoplasmic face of the Tat complex. This is followed by docking of the substrate to a second TatABC protomer. Then the positions of TatA and TatB are exchanged, and additional TatA copies are recruited to the translocase to enlarge the size of channel according to substrate. Driven by the proton motive force (*pmf*), the substrate is translocated across the membrane and released into the periplasm. During this process, the signal peptide is bound to the cytoplasmic side of TatC till the release of substrate (*26*). **b**, Under conditions with higher translocation throughput, more than one substrate can bind simultaneously to the Tat complex. One substrate undergoes docking, while the second binds only via its signal peptide. Similar to panel a, more copies of TatA are recruited, and the *pmf* drives the translocation of the docked substrate into the periplasm. The second substrate remains bound, then docks and queues for subsequent translocation.

Because *E. coli* has been used for a long time as a main model for Tat research, mutations of most key conserved residues have been performed and, as summarized in Table S4 and noted in the text above, the accumulated evidence fully supports our proposed mechanism. An alternative mechanism has been suggested in which the substrate is translocated through an independent oligomer of TatA (*10, 39*). While we cannot exclude this possibility, the mechanism by which the bound substrate would be transferred from the receptor TatABC complex to the TatA oligomer is unclear. Furthermore, if translocation occurs through an independent TatA oligomer, it is unclear why, according to our structure, the receptor Tat complex is apparently optimized to receive a substrate inside its cavity and undergo conformational changes (*14, 18, 35*– *37, 39*). It is however still feasible that only one of TatC interfaces opens up to accommodate many TatA subunits and so the translocation channel would be formed by the sideways-opened TatABC complex extended by extra TatA subunits (*20*). In summary, although the structure of the substrate-engaged TatBC complex is now clear, the precise organization of the TatA oligomers relative to the substrate-bound TatBC complex remains to be elucidated.

## Acknowledgments

We thank IST Austria EM facility for the use of Titan Krios TEM. Data processing was preformed using IST high-performance computer cluster. We thank Dr R. Roemhild and Professor C. Guet (ISTA) for help in constructing Tat deletion strains, and A. Charnagalov (ISTA) for technical help.

## Author contributions

Conceptualization: LAS Methodology: ZZ, LAS Investigation: ZZ, LAS Visualization: ZZ, LAS Funding acquisition: LAS Project administration: LAS Supervision: LAS

Writing – original draft: ZZ

Writing – review & editing: ZZ, LAS

## Data and materials availability

The cryo-EM map is deposited in the Electron Microscopy Data Bank under accession number EMD-53848. The model is deposited in the Protein Data Bank under accession number 9R91. Source data are provided within this paper.

## Supplementary Materials

Materials and Methods

Figs. S1 to S24

Tables S1 to S4

