## Supplementary Information for "Structure of *E. coli* Twin-arginine translocase (Tat) complex with bound cargo"

**The PDF file includes:**

Materials and Methods
Figs. S1 to S24
Tables S1 to S4

### Materials and Methods

#### Constructs and strains

Plasmids pBAD-TatABC, pBAD-ehTatABC, pBAD-TatABCstrep, pBAD-ehTatABCstrep, pBAD-TatABCflag and pBAD-TatABCflag were produced from pBAD plasmids. Natural sequence from TatA to TatC (with or without tag) was inserted between araBAD promoter and terminator. A 3XFlag sequence (GSDYKDHDGDYKDHDIDYKDDDDK) was added to the C-terminal end of TatC. An efficient ribosome binding site was added in front of TatA to enhance expression level of TatABC (*40*). Plasmids containing this configuration are designated with the prefix “pBAD-eh”. For overexpression of SufI, natural SufI fragments were inserted to pQlinkN plasmid (Addgene) (*41*). The ampicillin resistance gene was replaced by kanamycin resistance gene for compatibility with pBAD plasmids. The 8*His tag with eight histidine coding sequence was added to the C-terminus of SufI via primer overhang. The expression plasmids of TatABC and SufI were sequenced by Microsynth (https://srvweb.microsynth.ch/) to confirm correct sequences.

For deletion of TatABC from BL21 and Top10, a pSIM5 plasmid (obtained from NIH, USA)(*42*) was introduced to Top10 and BL21 cells. A DNA fragment containing an FRT-flanked kanamycin resistance gene was obtained from pKD4 (Addgene) by PCR reaction with additional 40 bp flanking regions homologous to the sequences upstream of *tatA* and downstream of *tatC* (*43*). This fragment was introduced to BL21/Top10 strains genome with pSIM5. After incubation at 42 °C cells were selected on kanamycin plates and PCR confirmed that *tatABC* fragment was deleted successfully. To excise the kanamycin resistance cassette, the pCP20 plasmid (*44*) was introduced into the knockout strains. Cultures were incubated at 42 °C to induce FLP recombinase expression and eliminate the kanamycin marker via FRT recombination. Colonies that lost kanamycin resistance were selected, and deletion of the marker was confirmed by PCR.

#### Functional tests

Tat substrates AmiA and AmiC were used as reporters to assess Tat functionality. AmiA and AmiC are N-acetylmuramoyl-L-alanine amidases involved in remodelling the cell wall during bacterial growth. In the absence of AmiA and AmiC, *E. coli* exhibits increased sensitivity to sodium dodecyl sulphate (SDS) (fig. S1) (*17, 45*). Overnight cultures of *E. coli* strains were diluted 1:25 in LB medium and grown to an $OD_{600}$ of approximately 0.9. Cultures were then normalized to an $OD_{600}$ of 1.0 before spotting onto LB agar plates containing 2% SDS. For comparison on plain LB plates, cells were diluted $1{:}10^5$ prior to inoculation. Plates were incubated overnight at 37 °C.

To test whether overexpressed SufI was translocated via the Tat pathway, immunoblotting was performed to detect both precursor and mature forms of SufI in various strains (fig. S2). Cells were grown under inducing conditions and then harvested. Cultures were normalized to an $OD_{600}$ of 3 using distilled water and mixed with SDS-PAGE loading buffer. Equal volumes of each sample were loaded onto an SDS-PAGE gel for electrophoretic separation. For immunoblot detection of TatC, an anti-Flag antibody (F1804, Sigma) was used as the primary antibody, followed by an HRP-conjugated anti-mouse IgG secondary antibody (NA9310V, Amersham ECL). For detection of SufI, an HRP-conjugated anti-His IgG antibody (1014992, QIAGEN) was used.

**Expression of Tat**

*E. coli* Top10 ΔTatABC strains harbouring the plasmids pBAD_ehTatABC-Flag and pQlinkN_SufI-His were initially cultured overnight (ON) in LB medium supplemented with ampicillin and kanamycin. The overnight culture was then diluted 1:50 into TB medium containing ampicillin and 0.05 mM arabinose to induce TatABC-Flag expression. After 3 hours of incubation at 37 °C with shaking at 200 rpm, 0.1 mM IPTG was added to induce SufI-His expression. Following an additional 3 hours of growth, cells were harvested by centrifugation at 8000 rpm at room temperature (RT). Cell pellets were flash-frozen in liquid nitrogen and stored at −80 °C for subsequent purification.

**Purification of TatBC-SufI**

Cell pellets were thawed at RT and resuspended in lysis buffer (50 mM Tris-HCl pH 7.6, 90 mM NaCl, 60 mM KCl, 0.5 M sucrose, 0.002% PMSF) to an $OD_{600}$ of 50, with protease inhibitor mix (Roche, EU) added. Lysozyme was added to 0.2 mg/mL, and the suspension was incubated for 30 min at RT. Re-crystallized digitonin was then added (25 mg/mL of cell suspension), followed by incubation at 4 °C for 1 hour. Lysates were centrifuged at 5000 × g for 15 min to remove debris. The supernatant was incubated with Anti-FLAG® M2 resin (Sigma, EU) at 50 µl per 1 ml culture for 1 hour at 4 °C. Resin was washed five times with buffer (50 mM Tris-HCl pH 7.6, 90 mM NaCl, 60 mM KCl, 20% glycerol, 0.2% digitonin, 0.002% PMSF), and proteins were eluted with the same buffer containing 0.3 mg/mL 3×FLAG peptide (F4799, Sigma).

Eluates were concentrated using a 100 kDa MWCO concentrator (Sigma) to <50 µl. To reduce glycerol concentration, an equal volume of buffer without glycerol was added, and the sample was reconcentrated. The sample was loaded onto a Superose™ 6 Increase 5/150 GL column (ÄKTA micro) and eluted with SEC buffer (50 mM Tris-HCl pH 7.6, 90 mM NaCl, 60 mM KCl, 10% glycerol, 0.2% digitonin, 0.002% PMSF). Elution profiles are shown in fig. S3a. Tat-containing fractions were pooled and incubated overnight at 4 °C with 100 µl nickel resin (fig. S3bc). Protein was eluted with SEC buffer (without glycerol) containing 300 mM imidazole, then concentrated, diluted with SEC buffer (without glycerol) to reduce imidazole concentration and reconcentrated for downstream use. After this step the preparation was highly enriched in SufI content with concomitant partial loss of TatA (fig. S3b).

**Grids preparation**

Quantifoil 0.6/1 300 mesh copper grids were manually coated (Leica ACE600) with an approximately 0.9 nm layer of continuous carbon and glow discharged just before usage for 5 sec at 25 mA. 3 µl of fresh sample (0.25 mg/ml) was applied to the grids in a humidified chamber (100% humidity at 4 °C) of a Vitrobot Mark IV, blotted for 2 s with 25 force and plunge-frozen in liquid ethane.

**Data collection**

The micrographs were collected on a 300 kV Titian Krios G3i, equipped with Gatan K3 direct detector, in super-resolution mode. 11,284 movies were collected in 48 hours using EPU. The data collection details are summarized in Table S1.

**Processing**

The processing pipeline is illustrated in fig. S4. The micrographs were processed using Relion 5.0 (*46*). MotionCor2(*47*) was used for motion correction and CTFFIND4 (*48*) for the estimation of micrographs' CTF parameters. After motion correction and CTF estimation, 10,177 good micrographs were selected. Particles were first picked with default Topaz model (*49*) in Relion using a subset of 400 micrographs. After 2D classification, good classes were select as references for Topaz training and picking on all micrographs. After 2D classification of all particles, 1.3 M particles in good classes (showing Tat complex with attached SufI in one or two copies) were selected and subjected to 3D classification. Many different 3D classification strategies with different masks were attempted in order to identify the best-defined classes. Classifications with alignments did not reveal clear differences between classes, suggesting relatively uniform conformation of particles. Consistently, 3D refinement of all particles with mask containing TatBC and SufI led to an averaged map of about 4 Å resolution. The best class was identified by subjecting thus refined particles to 3D classification without alignment into 5 classes with regularization parameter T=64. The best class with 164k particles was selected for further global and local 3D refinements, CTF refinement, and Bayesian polishing. Subsequently, the micelle density was removed using particle subtraction with mask around TatBC-SufI, followed by another round of local 3D refinement and post-processing. This refinement yielded the final map at a gold-standard resolution of 3.74 Å. Local resolution was estimated and map filtered by Relion. The core of TatBC is at about 3.5 resolution, while SufI is less ordered, particularly at its distal tip. Local refinement with mask covering only SufI did not improve its resolution.

Two-SufI particles were selected and 3D refined with masks including micelle and either one or two SufI molecules. Second SufI usually showed elongated density due to its flexible position (fig. S6b right). 3D classification with alignment separates a two-SufI class with 140k particles. 3D refinement of this class lead to relatively well-defined second SufI but overall resolution remained low (fig. S6b left). 300k two-SufI particles from 2D classification were also 3D refined with mask including micelle, Tat and 1 SufI, and then separated by 3D classification into 5 classes without alignment using T=64. 4 out of 5 classes show an additional helix-like density, likely of bound second signal peptide (fig. S24).

**Model building and bioinformatics**

Aphafold3 (https://alphafoldserver.com/) (*50*) structure of 3 TatC, 3 TatB, 3 TatA and 3 SufI, which correctly places signal peptide (but does not put the rest of SufI on the adjacent TatBC unit), was used as initial model after deleting regions not present in EM map. The model was first rigid body fitted into the cryo-EM density, followed by manual adjustments in COOT.

Reference model and secondary structure restrains were used during phenix.real_space_refine runs (*51*) to keep correct geometry of Tat in less defined regions. The refined structure was then re-build in COOT with cycles of phenix.real_space_refine. The bulk of SufI was refined as a rigid body, apart from better defined areas at the interface with Tat. The final structure was validated by Molprobity (*52*). Refinement statistics are shown in Table S1. Structure figures were made using ChimeraX (*53*). The conservation analysis of Tat was done using ConSurf server at: (https://consurf.tau.ac.il/) (*54*). Electrostatic solvent-accessible surface potentials were calculated using the PyMOL APBS electrostatics Plugin (*55*) with default template settings. Electrostatic solvent-excluded surface potentials were calculated in ChimeraX using "coulombic" command.

### Methods References

50. J. Abramson, J. Adler, J. Dunger, R. Evans, T. Green, A. Pritzel, O. Ronneberger, L. Willmore, A. J. Ballard, J. Bambrick, S. W. Bodenstein, D. A. Evans, C.-C. Hung, M. O'Neill, D. Reiman, K. Tunyasuvunakool, Z. Wu, A. Žemgulytė, E. Arvaniti, C. Beattie, O. Bertolli, A. Bridgland, A. Cherepanov, M. Congreve, A. I. Cowen-Rivers, A. Cowie, M. Figurnov, F. B. Fuchs, H. Gladman, R. Jain, Y. A. Khan, C. M. R. Low, K. Perlin, A. Potapenko, P. Savy, S. Singh, A. Stecula, A. Thillaisundaram, C. Tong, S. Yakneen, E. D. Zhong, M. Zielinski, A. Žídek, V. Bapst, P. Kohli, M. Jaderberg, D. Hassabis, J. M. Jumper, Accurate structure prediction of biomolecular interactions with AlphaFold 3. *Nature* **630**, 493–500 (2024).

a

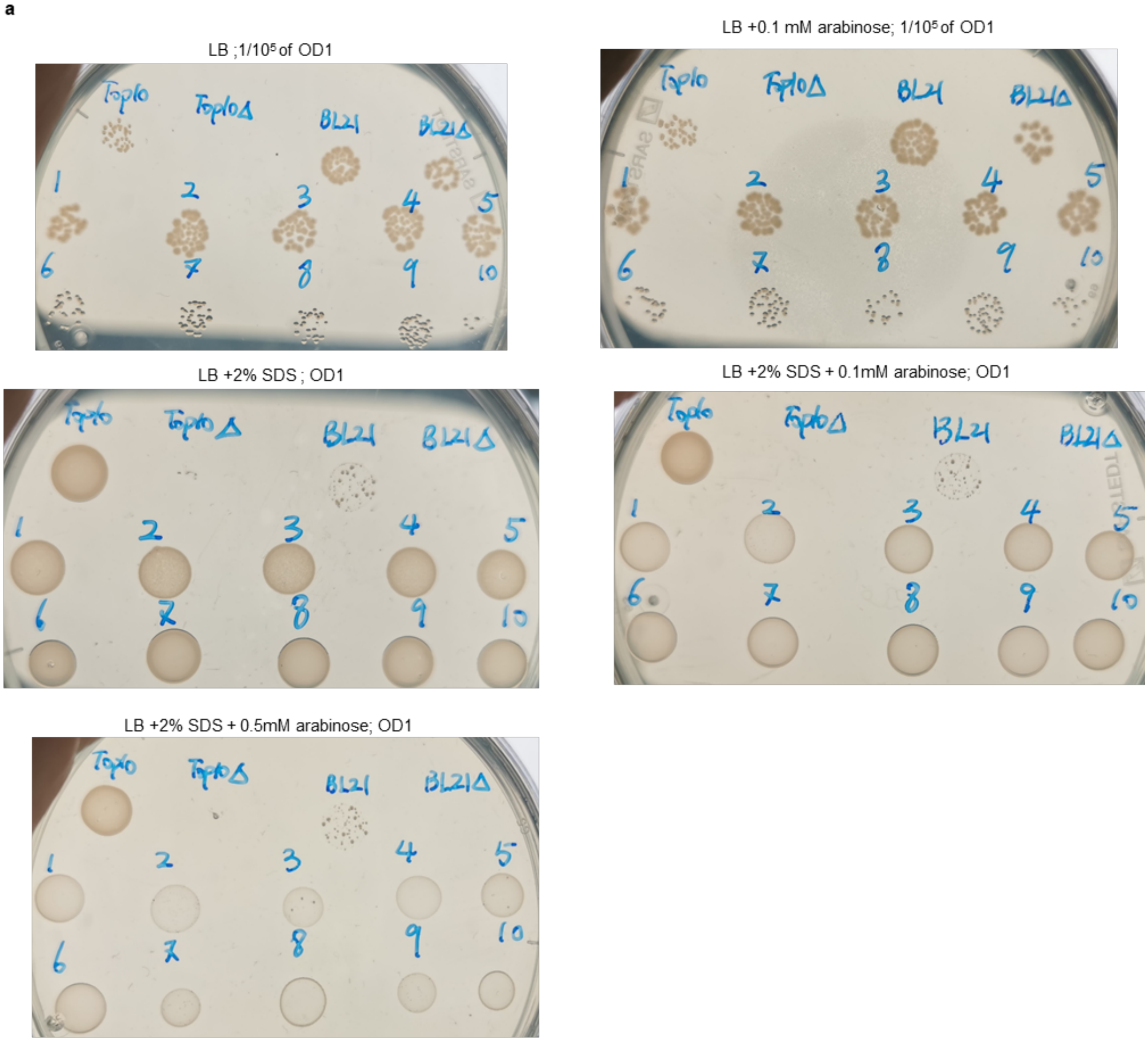


b

| | | | | | |
|---|---|---|---|---|---|
| | Top10; | Top10Δ; | BL21; | BL21Δ | |
| **BL21Δ +**: | 1.pBAD-TatABC-strep; | 2.pBAD-ehTatABC-strep; | 3.pBAD-TatABC-Flag; | 4.pBAD-ehTatABC-Flag; | 5.pBAD-TatABC |
| **Top10Δ +**: | 6.pBAD-TatABC-strep; | 7.pBAD-ehTatABC-strep; | 8.pBAD-TatABC-Flag; | 9.pBAD-ehTatABC-Flag; | 10.pBAD-TatABC |

**Fig. S1. Phenotypes of *E. coli* strains.**

Overnight cultures of *E. coli* strains (listed in **b**) were diluted 1:25 and grown to an $OD_{600}$ of ~0.9. Cultures were normalized to $OD_{600} = 1$ and spotted onto LB plates containing 2% SDS. For comparison on plain LB plates, cells were diluted $1{:}10^5$ times before spotting. Plates were incubated overnight at 37 °C. Growth conditions are indicated above each panel. **b**, Table of *E. coli* strains and corresponding plasmids. "eh-" prefix represents plasmids containing efficient ribosome binding site that enhances expression level. The top row of the plate corresponds to either wild-type strains (Top10/BL21) or *tatABC* deletion mutants (Top10Δ/BL21Δ). The second and third row indicate different pBAD plasmids with BL21Δ and Top10Δ, respectively. Deletion of *tatABC* in Top10 leads to slower growth.

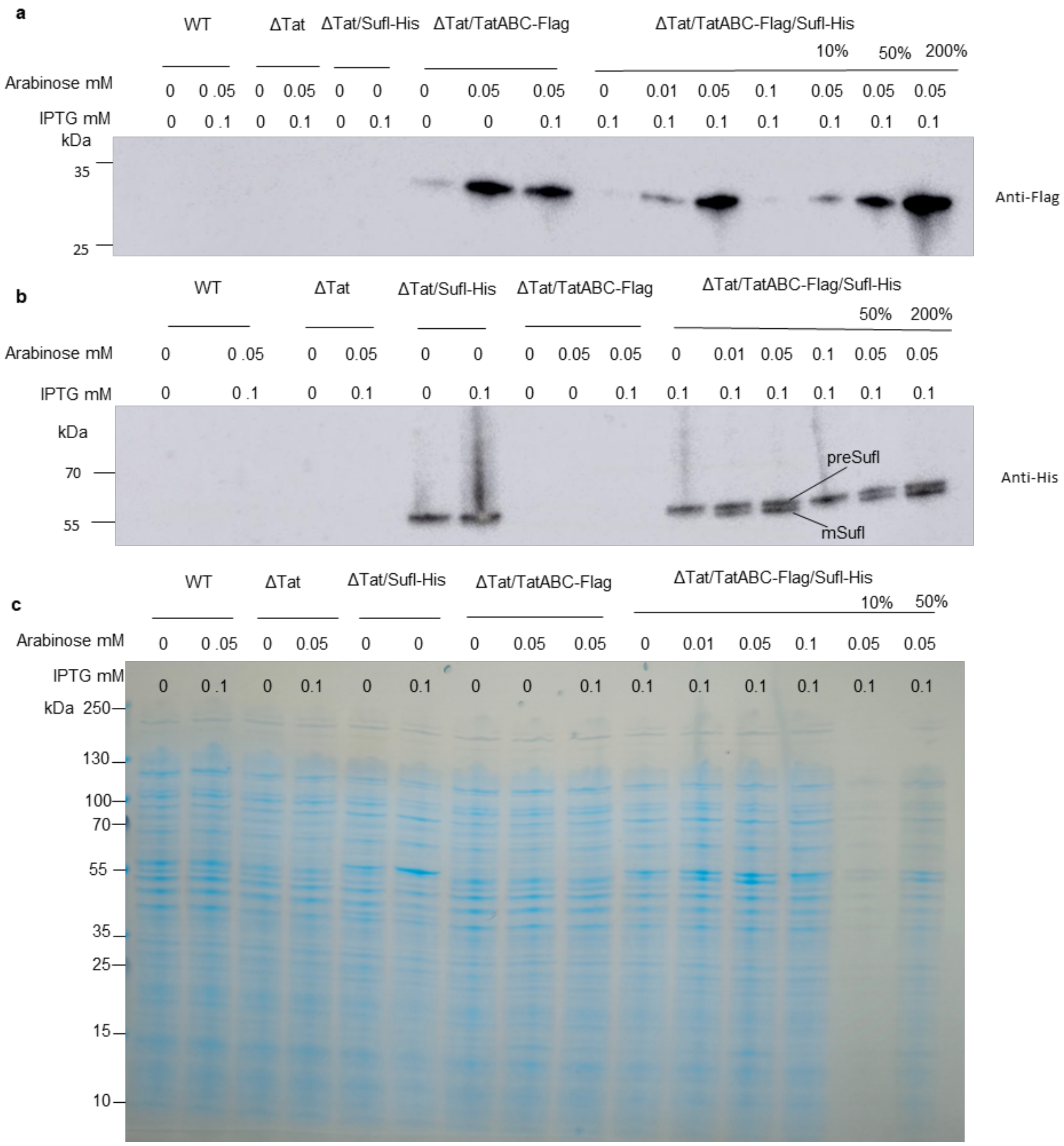


**Fig. S2. SDS-PAGE with immunoblot and Coomassie staining testing activity of Tat translocating SufI.**

**a,** Anti-Flag immunoblot of different Top10 *E. coli* strains with different concentrations of arabinose to detect TatC expression. **b,** Anti-His immunoblot to estimate pre-SufI and mature SufI levels (expression induced by IPTG) to assess Tat translocation activity. **c,** Coomassie staining of samples used for immunoblot.

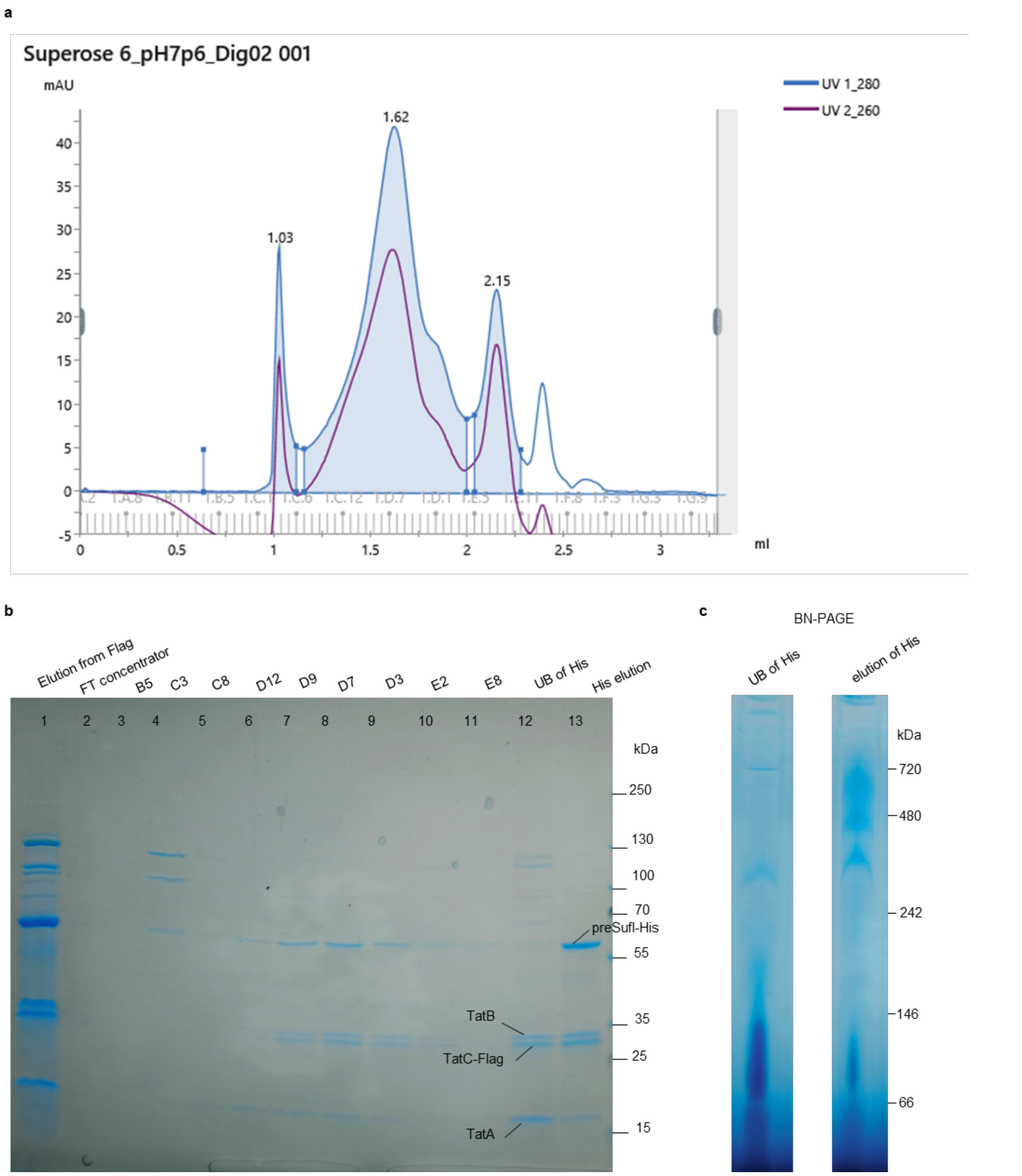


**Fig. S3. Purification of TatBC/SufI from Top10 ΔTat/TatABC-Flag/SufI-His strain.**

**a**. Size exclusion chromatography profile of harvested protein from Flag pull-down (lane 1 in b). **b**. SDS-PAGE and Coomassie staining of protein fractions during purification. Lane 1 represents concentrated protein harvested from Flag pull-down. Lane 2 represents protein flow through 100 kDa cut-off protein concentrator. Lanes 3-11 represent fractions of SEC. Lane 12 represents fractions not bound to nickel resin. Lane 13 represents elution from nickel resin. **c**. BN-PAGE and Coomassie staining of fractions not bound to nickel resin (UB of His) and eluted from nickel resin (elution of His).

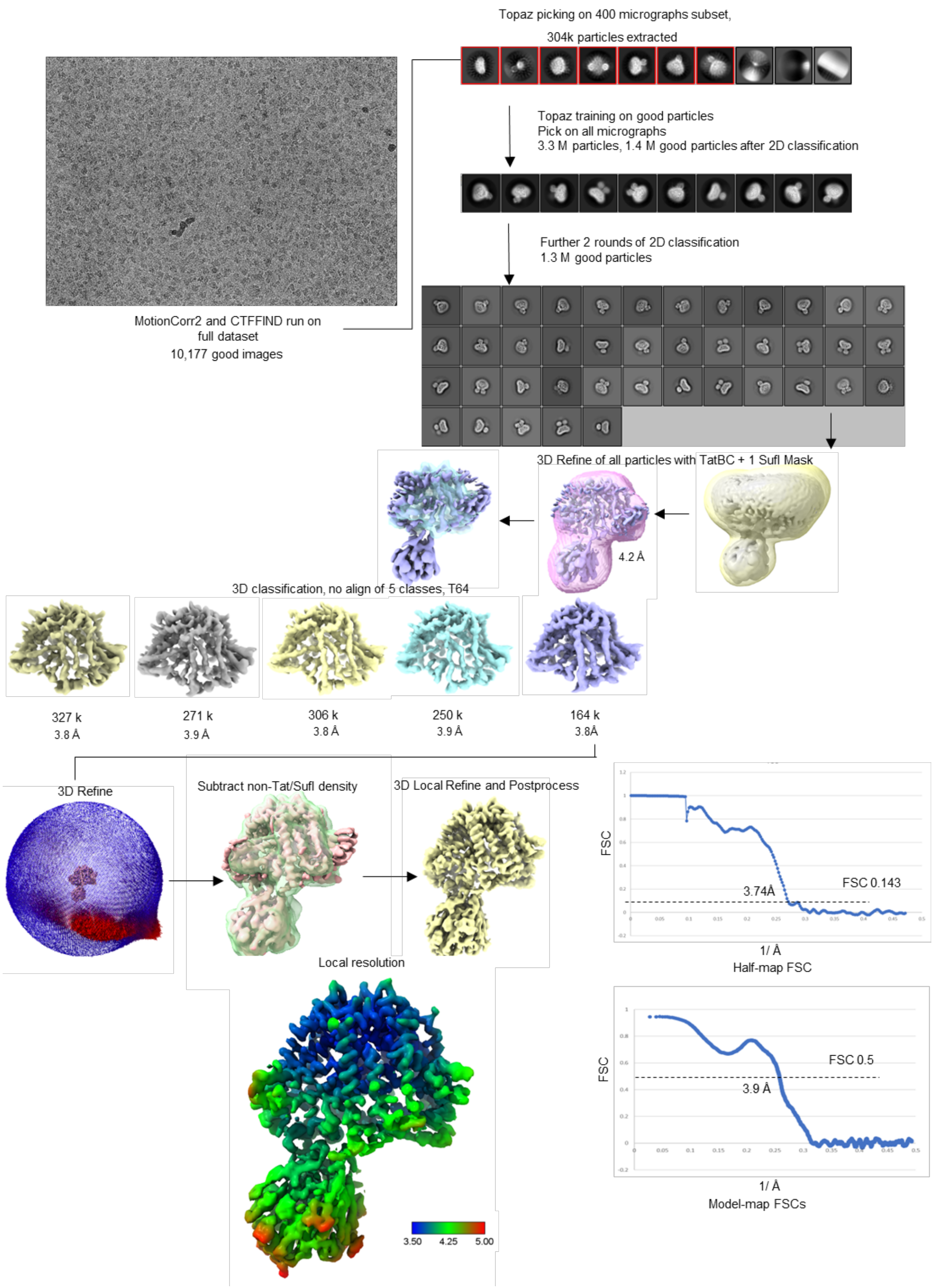


**Fig. S4. Cryo-EM data processing workflow.**

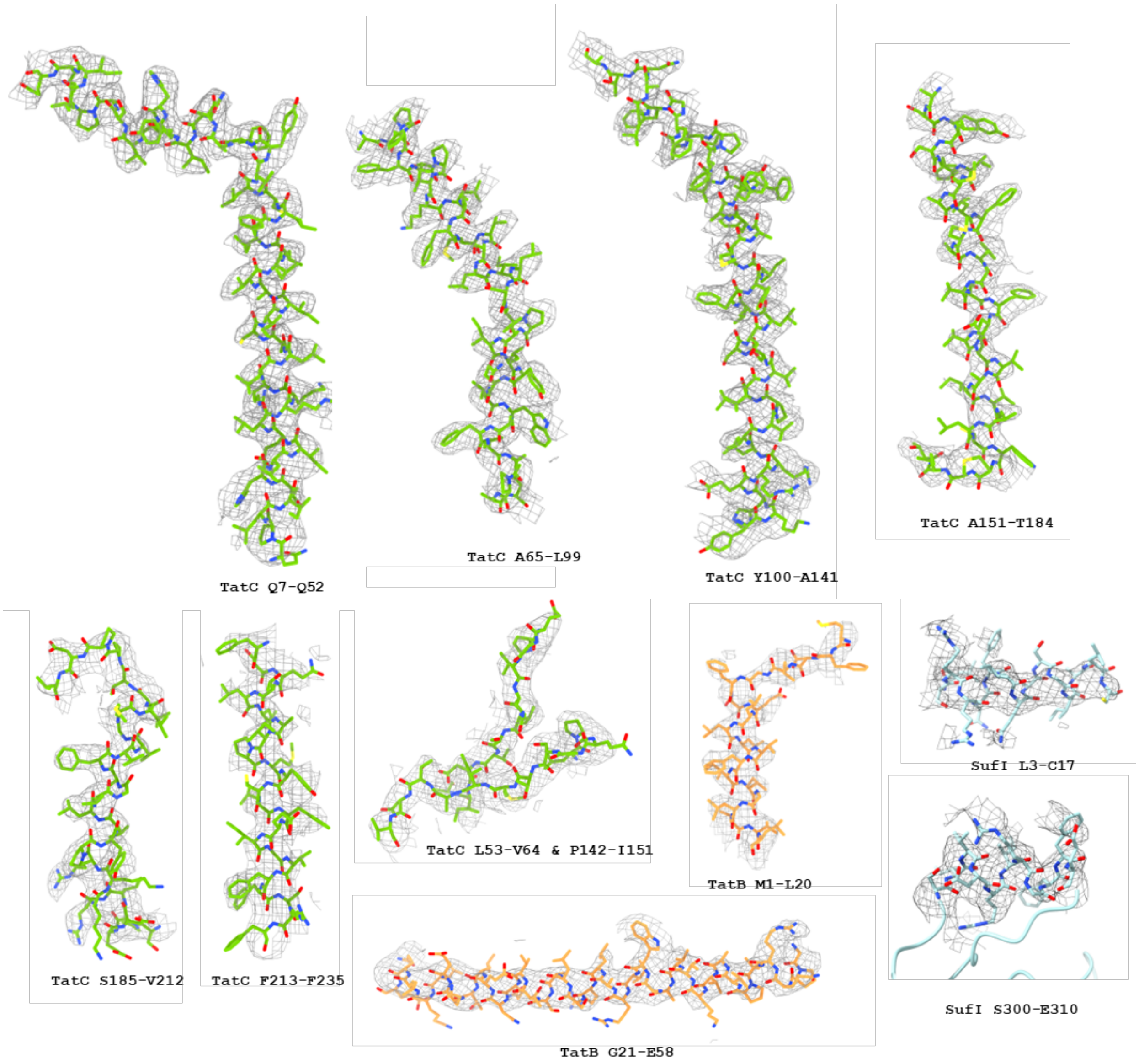


**Fig. S5. Examples of cryo-EM density fitting of Tat complex subunits.**

Cryo-EM density (local resolution-filtered in Relion 5.0) of $TatC_{amp\_helix}$, $TatB_{signal}$, signal peptide and amphipathic helix of SufI.

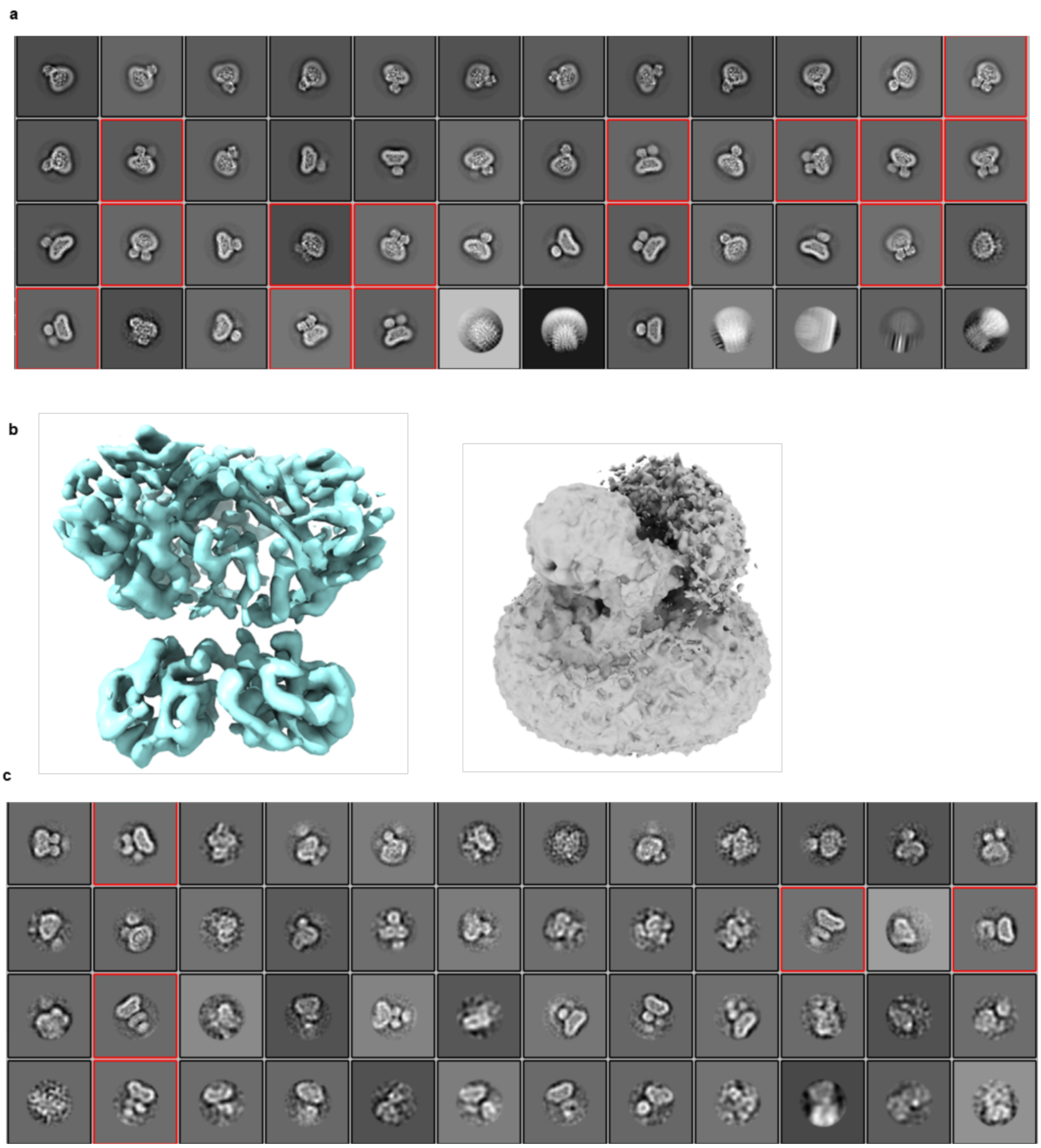

**Fig. S6. 2D Classification of all particles and 3D maps of TatBC-two-SufI Particles.**

**a**, Final 2D classification of TatBC/SufI Particles. Two-SufI 2D classes are highlighted, representing ~358k out of 1.32 M total good particles identified during 2D classification, corresponding to ~27% of the dataset. **b**, left, side-view of cryo-EM map of 140k two-SufI particles 3D refined with mask containing micelle+2 SufI (best class from 3D classification with alignment of all particles). The right panel shows a cytoplasmic view of the cryo-EM map of 300k two-SufI particles refined with mask including micelle+1SufI, showing elongated fuzzy density at the second SufI position. Micelle density remains strong in both cases. **c**, One of 2D classifications of "not standard" TatBC/SufI particles. 2D class averages showing crown-shaped or three SufI-like densities at the cytoplasmic side of Tat are highlighted. These classes contain ~1.3 k particles out of 1.2 M total good particles from this round of classification, representing ~0.1% of the dataset.

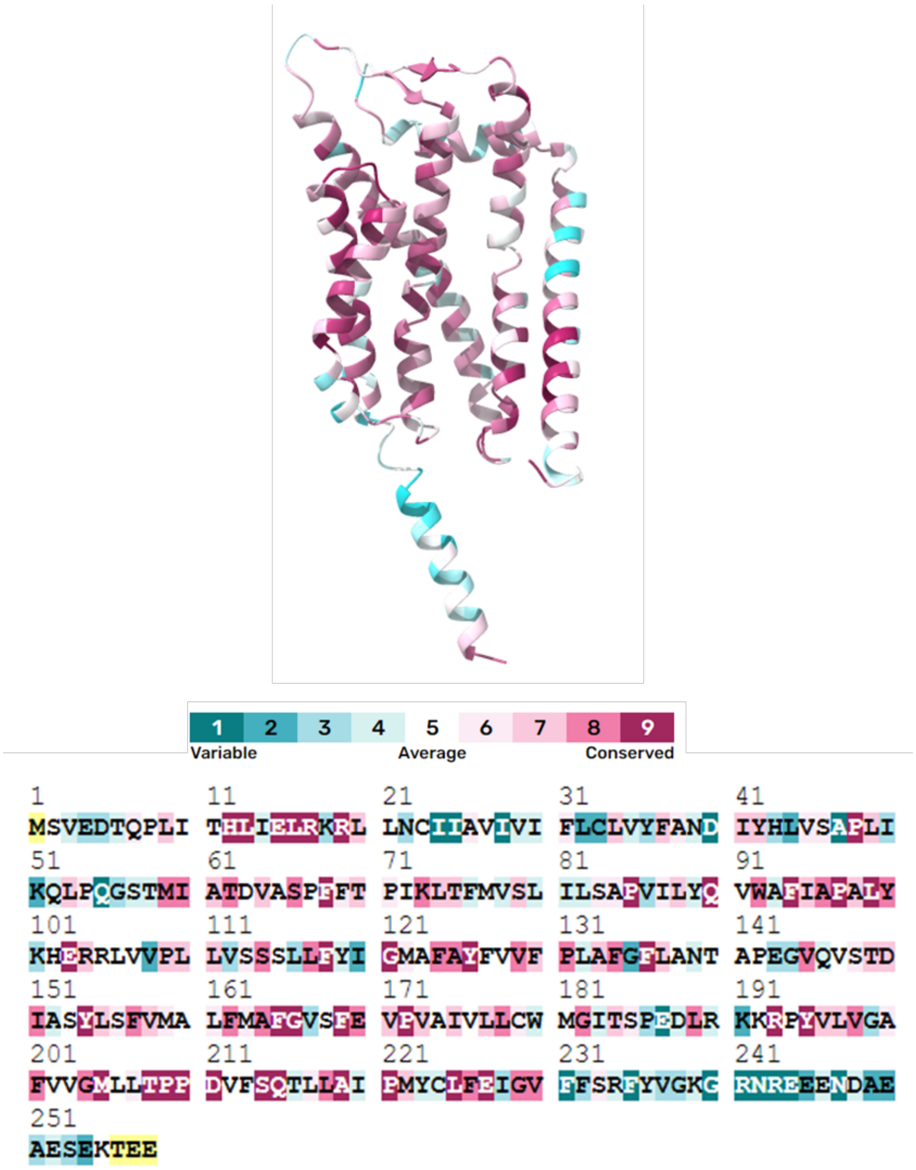


**Fig. S7. Conservation scores of full TatC sequence.**

Residues of TatC were scored by conservation using Consurf server (https://consurf.tau.ac.il/ ). Residues with high conservation are colored maroon and low conservation cyan, displayed on AF3 model. Residues with unreliable conservation scores are colored yellow.

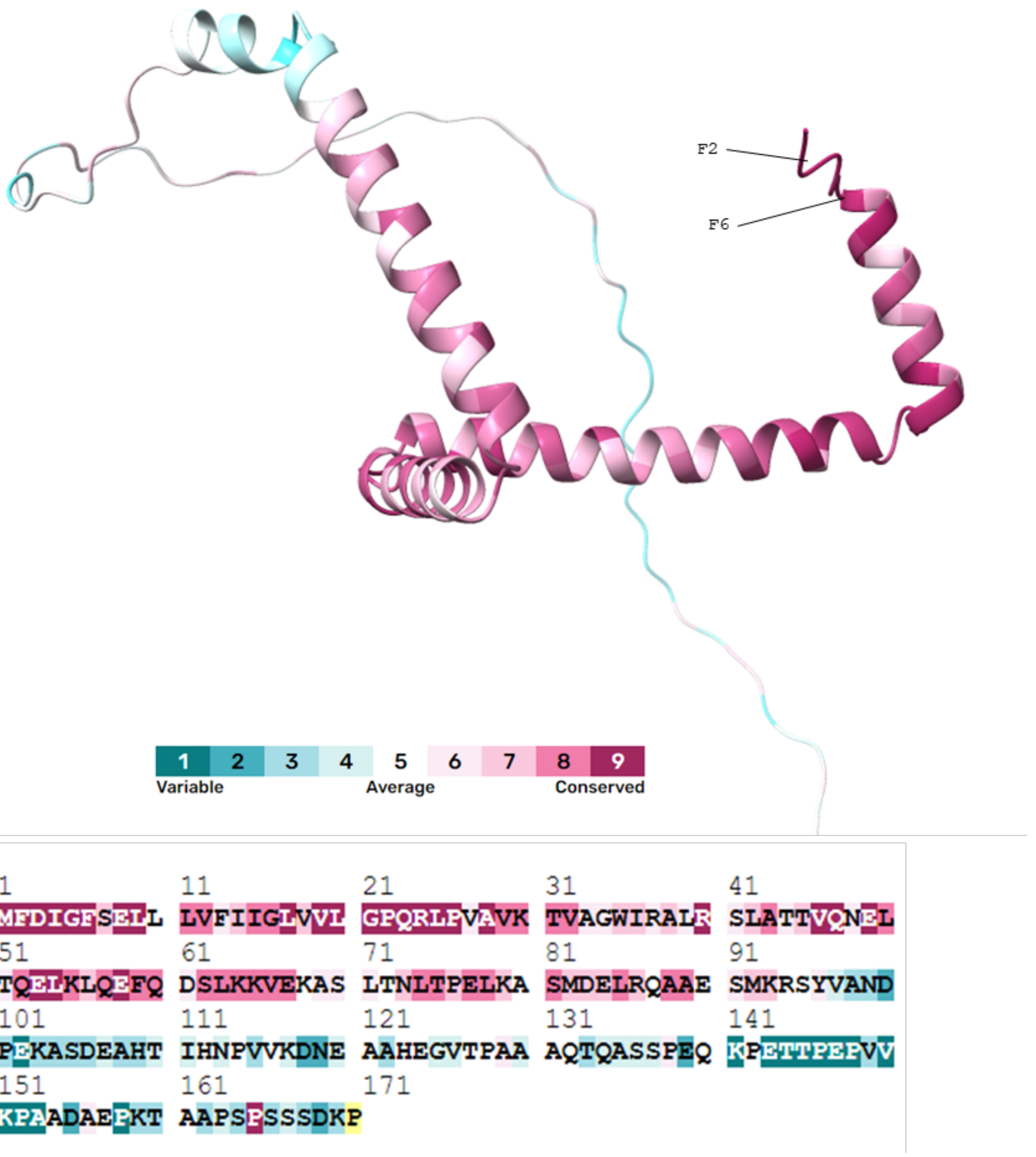


**Fig. S8. Conservation scores of full TatB sequence.**

Residues of TatB were scored by conservation using Consurf server (https://consurf.tau.ac.il/ ). Residues with high conservation are colored maroon and low conservation cyan, displayed on AF3 model. Residues with unreliable conservation scores are colored yellow.

a

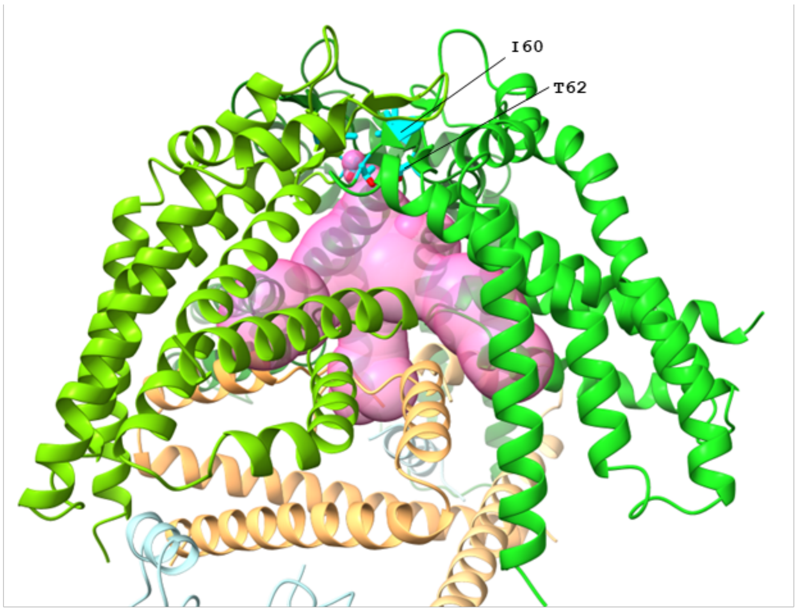


b

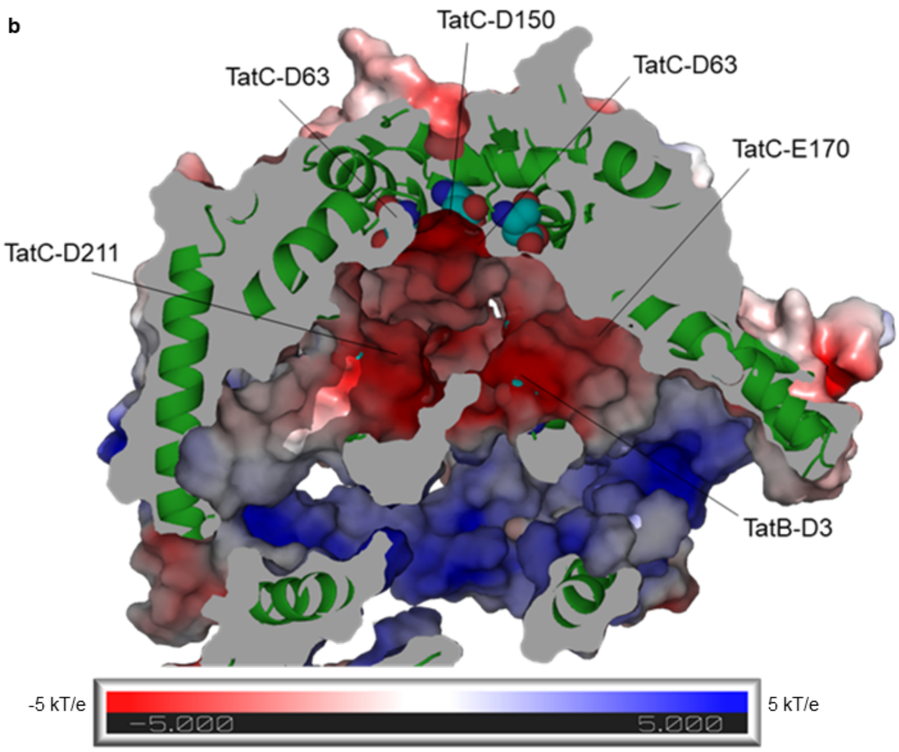


**Fig. S9. The channel and electrostatic surface potential of inner cavity of TatBC complex.**

**a,** Central cavity in TatBC complex is sealed by TatC Ile60 on the periplasmic side. The central cavity (calculated in MOLEonline server https://mole.upol.cz/online/) is shown in hot pink and I60 and T62 are shown in cyan as sticks. The constricted part of the pore around Ile60 has 0.9-1.1 Å radius, not allowing solvent (1.4 Å) to pass. The periplasm is at the top. **b,** Tat residues contributing to the negative charge of the inner cavity. Electrostatic solvent-accessible surface potential was calculated using the PyMOL APBS electrostatics Plugin with default template settings.

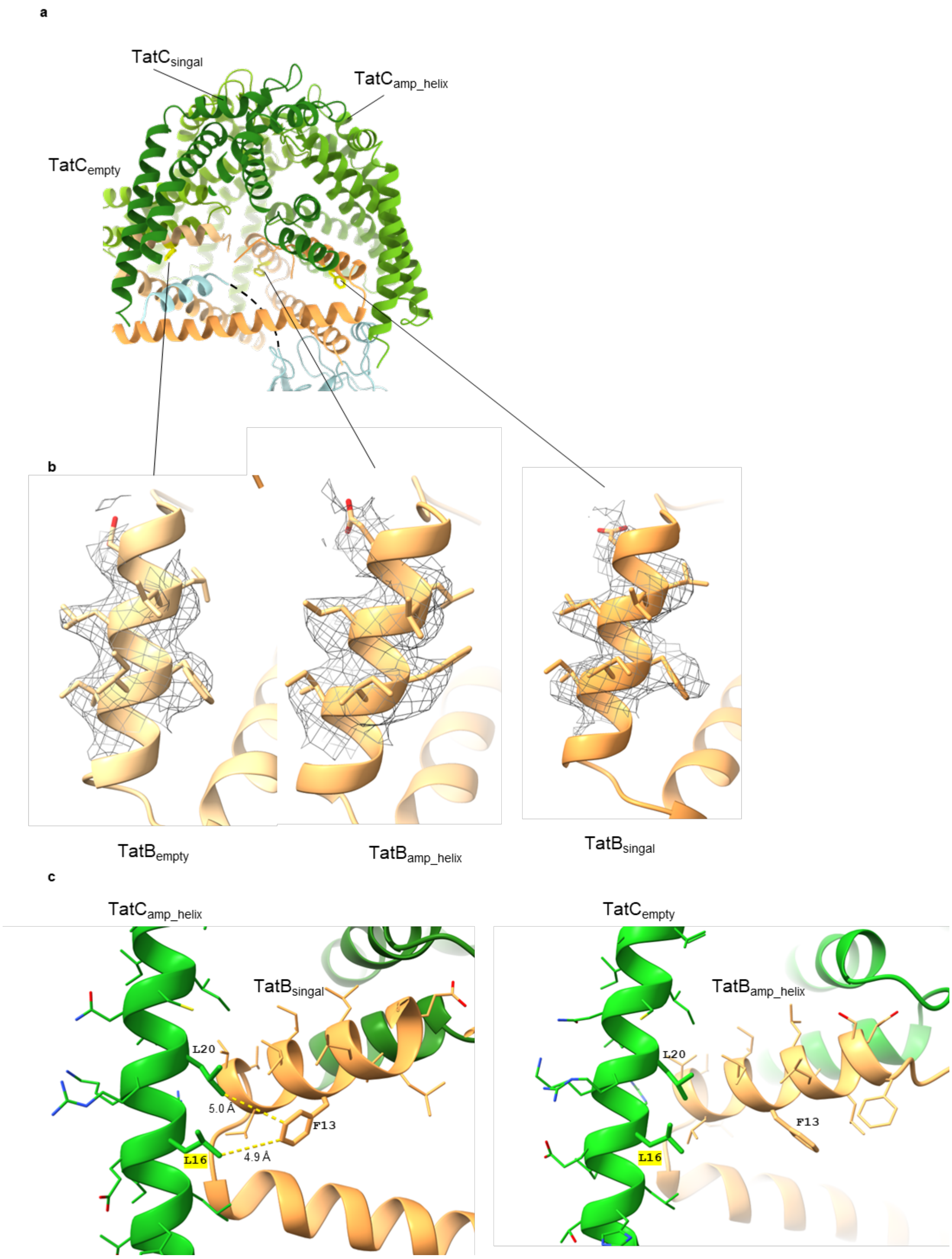


**Fig. S10. Cryo-EM density and conformation of TatB F13.**

**a**, Overview of TatB position in TatBC complex. **b**, Density fitting of side chains of TatB-F13. **c**, Hydrophobic interactions around $TatB_{signal}$-F13 (left) and $TatB_{amp\_helix}$-F13 (right). Hydrophobic interaction is shown in yellow dash lines. Conserved residues are highlighted.

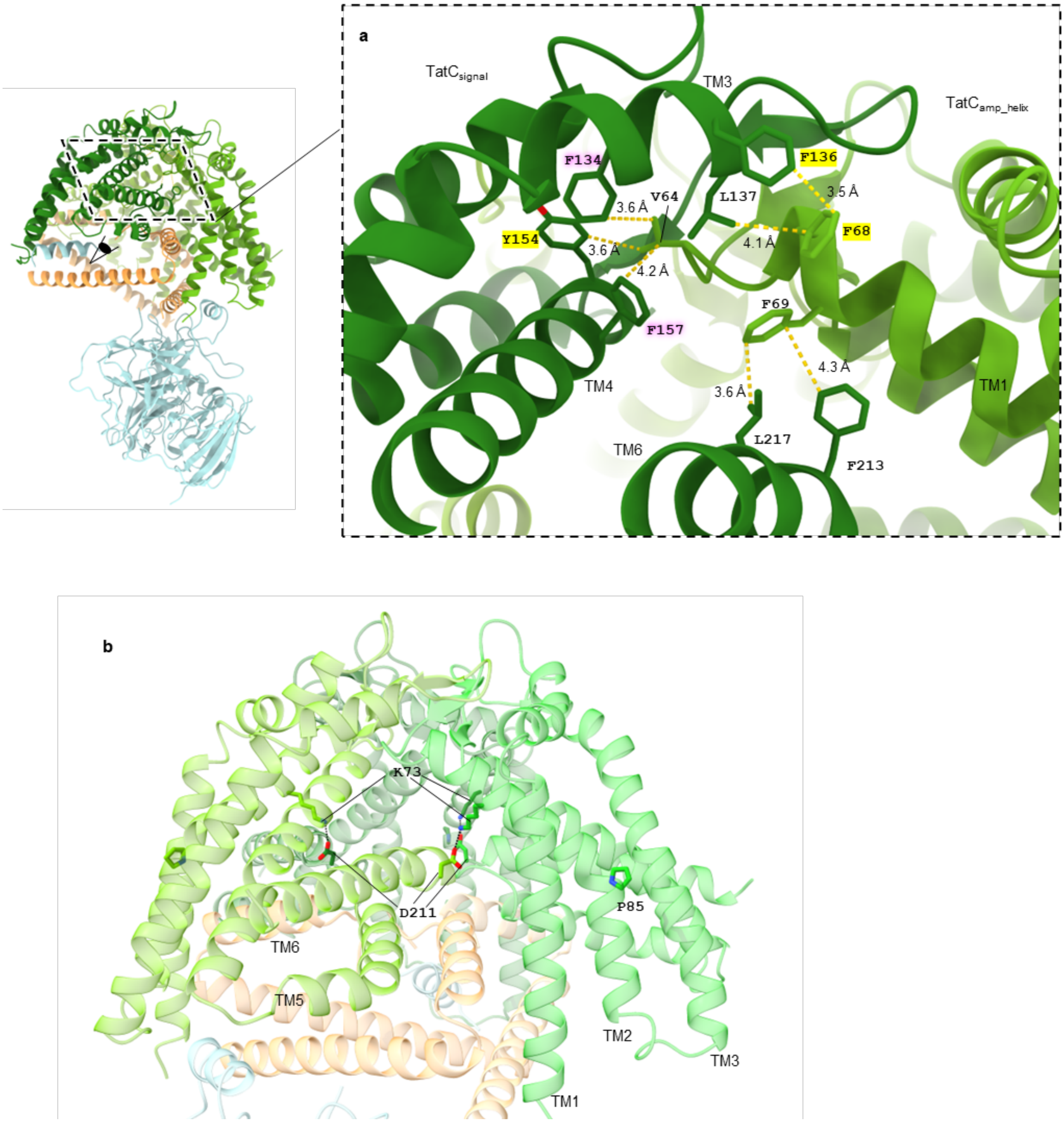


**Fig. S11. Hydrophobic interactions and salt bridges at the interface of two TatC monomers.**

**a**, Hydrophobic interactions between TatC$_{signal}$ and TatC$_{amp_helix}$ are shown. TM1 of TatC$_{amp_helix}$ interacts with TM3, TM4 and TM6 of TatC$_{signal}$. Specifically, V64 from TM1 interacts with F134 on TM3, and Y154 and F157 on TM4 of the neighbouring TatC. F68 from TM1 engages with F136 and L137 from TM3, while F69 interacts with F213 and L217 on TM6 of the adjacent TatC. Highly conserved residues (ConSurf score 9) are highlighted in yellow; conserved residues (ConSurf score 8) are shaded in pink. **b**, Salt bridges between TatC K73 and adjacent TatC D211. TatC K73 locates on TM2 that is bent by P85. Tat subunits are shown in transparent cartoons for clarity.

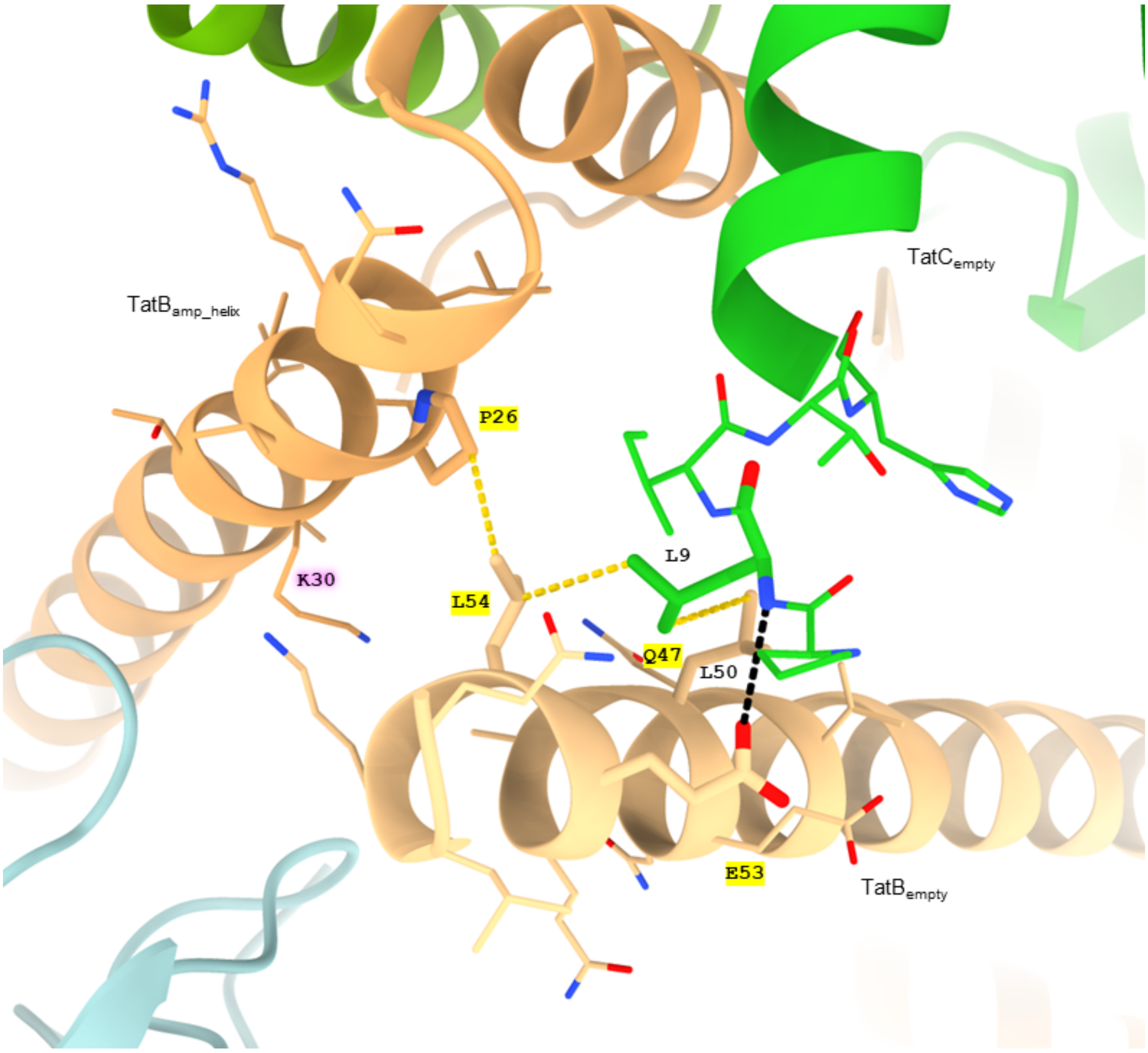


**Fig. S12. Interactions between the C-terminal end of the TatB amphipathic helix and the N-terminal end of TatC, along with N-terminal end of the amphipathic helix of adjacent TatB.**

A putative H-bond is indicated by a black dashed line, and potential hydrophobic interactions are shown as yellow dashed lines. Due to limited local resolution in this region, interaction distances are not displayed. A possible H-bond may form between the side chain of TatB E53 and the backbone of TatC L9. The hydrophobic side chain of TatC L9 is positioned near TatB residues L50 and L54, suggesting potential hydrophobic interactions. TatB L54 locates close to adjacent TatB P26, and TatB Q47 is close to K30 from adjacent TatB. Highly conserved residues (ConSurf score 9) are highlighted in yellow, and conserved residues (ConSurf score 8) are shaded in pink.

.

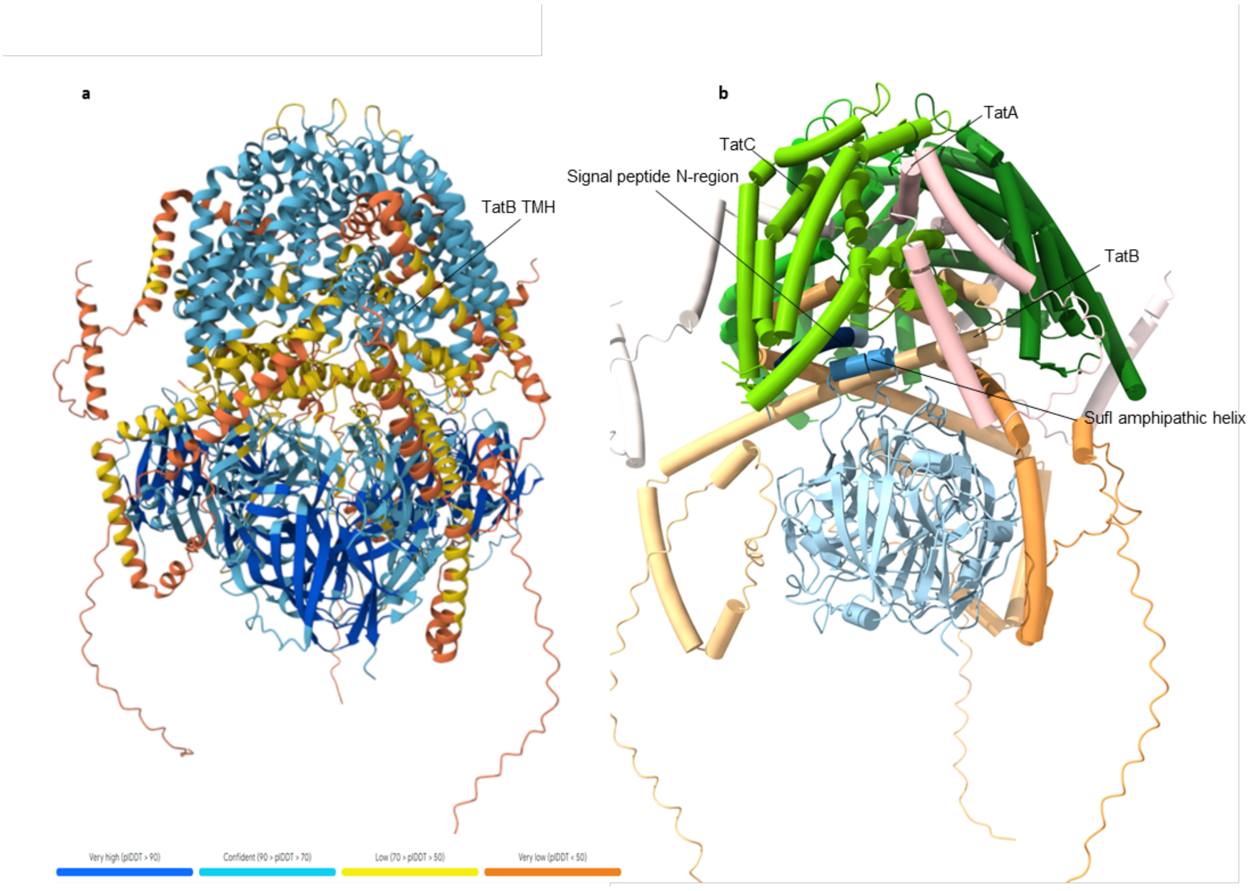


**Fig. S13. AF3-predicted model of TatABC trimer with three copies of SufI.**

Three copies each of *E. coli* TatA, TatB, TatC and pre-SufI were used as input. **a**, Per-atom confidence (pLDDT) of the AlphaFold prediction is color-coded from high (blue) to low (yellow to orange). **b**, Complex shown as cartoon cylinders colored by subunit. The signal peptide is shown in dark blue, and the short amphipathic helix SA is shown in blue. Only one SufI is shown, with the other two SufI omitted for clarity.

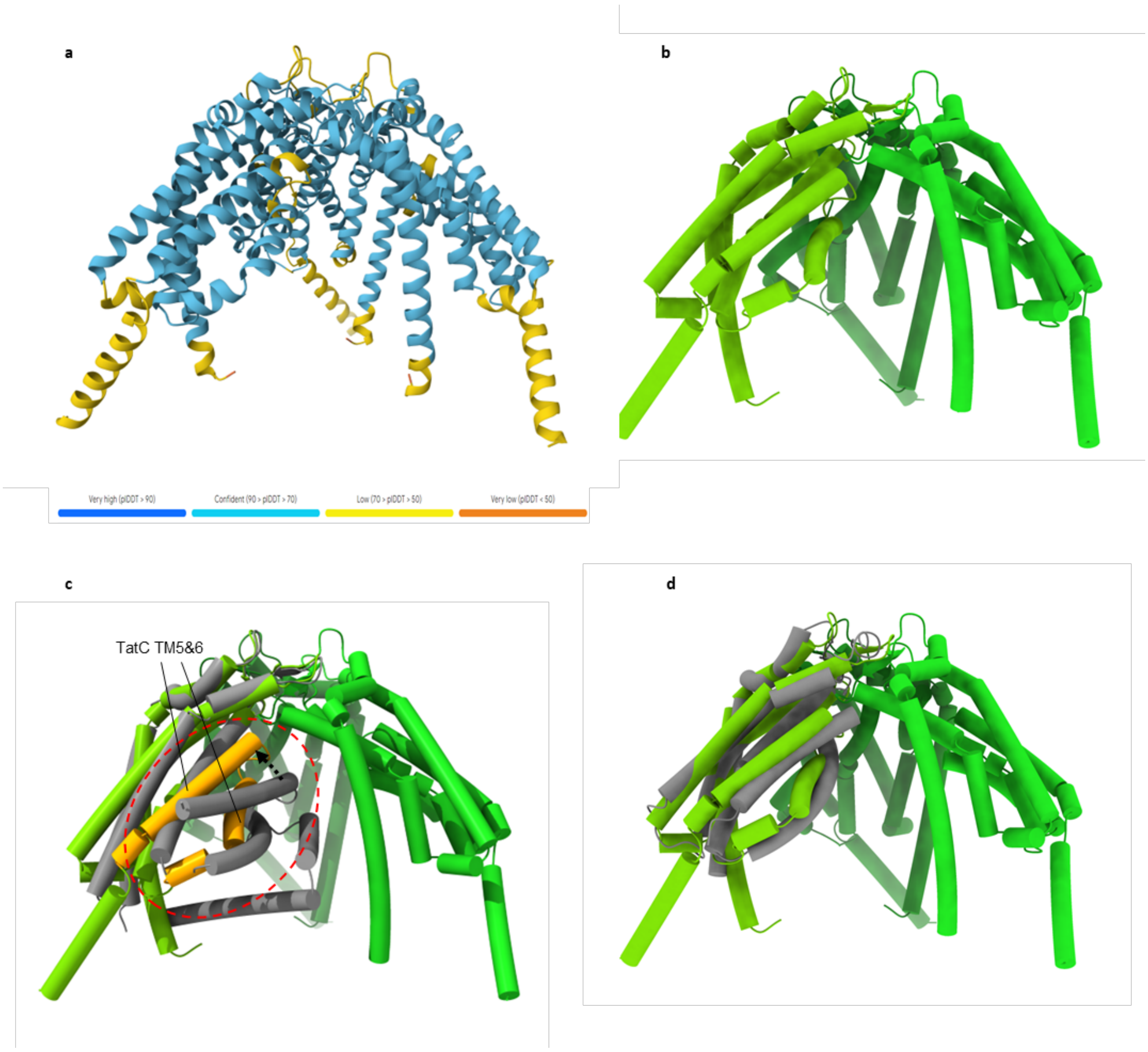


**Fig. S14. AF3-predicted model of $TatC_3$.**

Three copies of *E. coli* TatC were used as an input. **a**, Per-atom confidence (pLDDT) of the AlphaFold prediction is color-coded from high (blue) to low (yellow to orange). **b**, $TatC_3$ complex shown as cartoon cylinders. **c**, Structure comparison between AF3 $TatC_3$ (green) and our $TatB_3C_3$-SufI complex (grey). Only one copy of TatBC unit is shown for clarity. The TM5-6 from $TatC_3$ is highlighted in orange. **d**, Superposition of $TatC_3$ and *Aquifex* TatC crystal structure (grey, PDB ID: 4B4A).

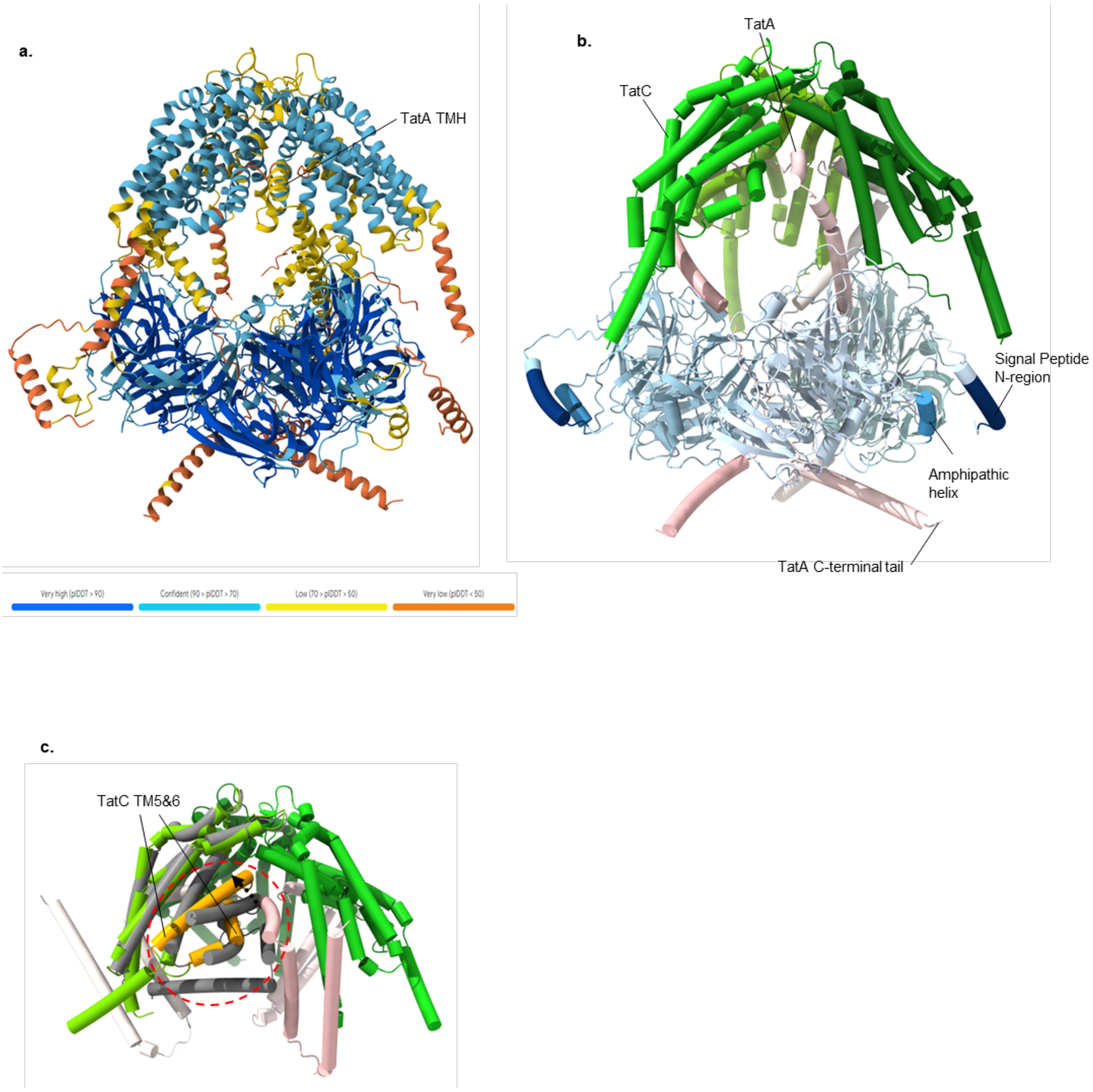


**Fig. S15. AF3-predicted model of the TatAC trimer bound to three copies of SufI.**
Three copies each of *E. coli* TatA, TatC, and pre-SufI were used as input. **a**, Per-atom confidence (pLDDT) of the AlphaFold prediction is color-coded from high (blue) to low (yellow to orange). **b**, Complex shown as cartoon cylinders colored by subunit. The signal peptide is shown in dark blue, and the short amphipathic helix is shown in blue. **c**, Structural alignment of the TatAC trimer (colored) with the TatBC trimer (grey), superimposed via one TatC subunit, reveals a looser architecture of the TatAC complex. TatC TM5 and 6 are colored in orange in $TatA_3C_3$ complex.

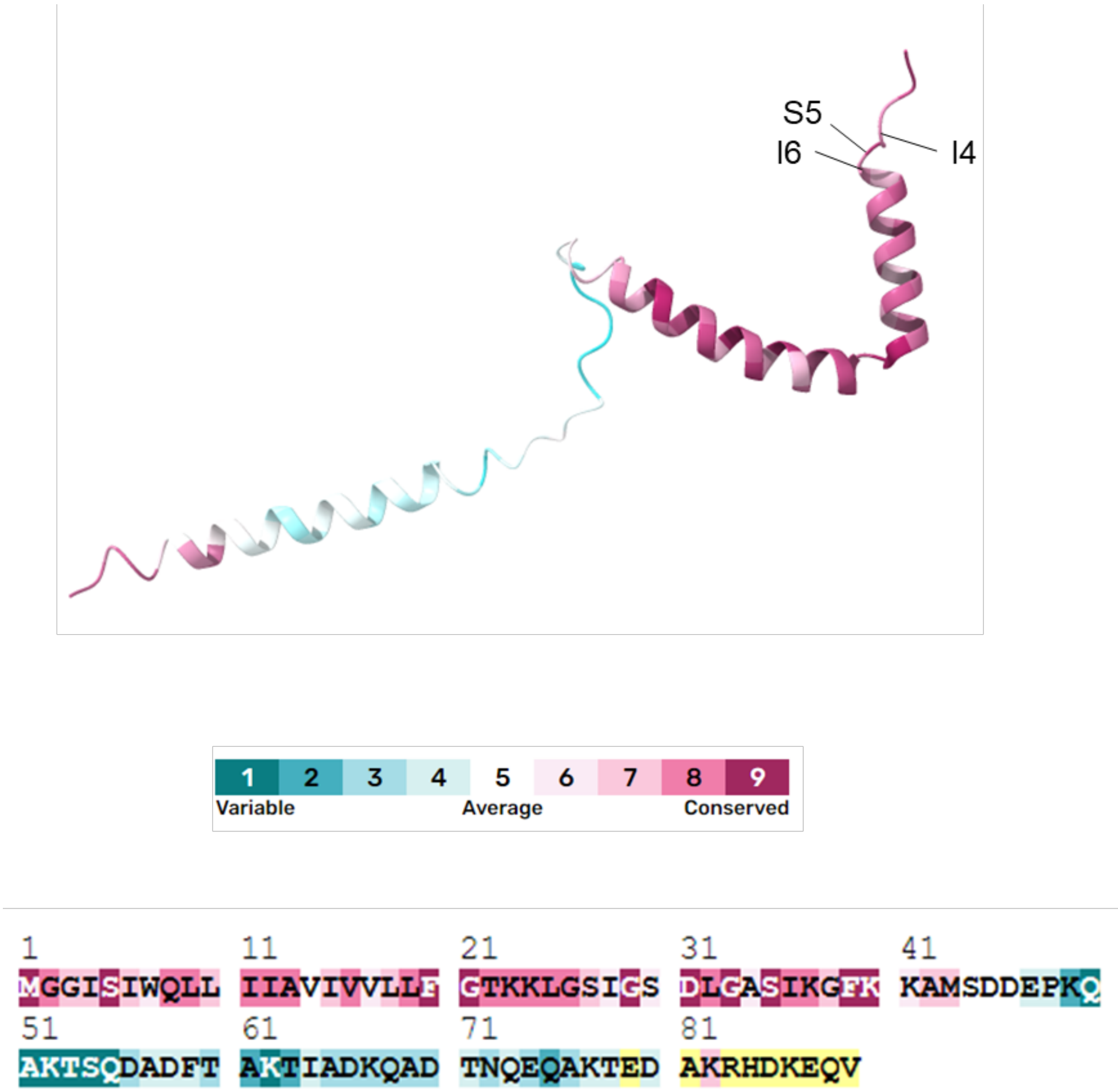


**Fig. S16. Conservation scores of full TatA sequence.**

Residues of TatA were scored by conservation using Consurf server (https://consurf.tau.ac.il/ ). Residues with high conservation are colored maroon and low conservation cyan, displayed on AF3 model. Residues with unreliable conservation scores are colored yellow.

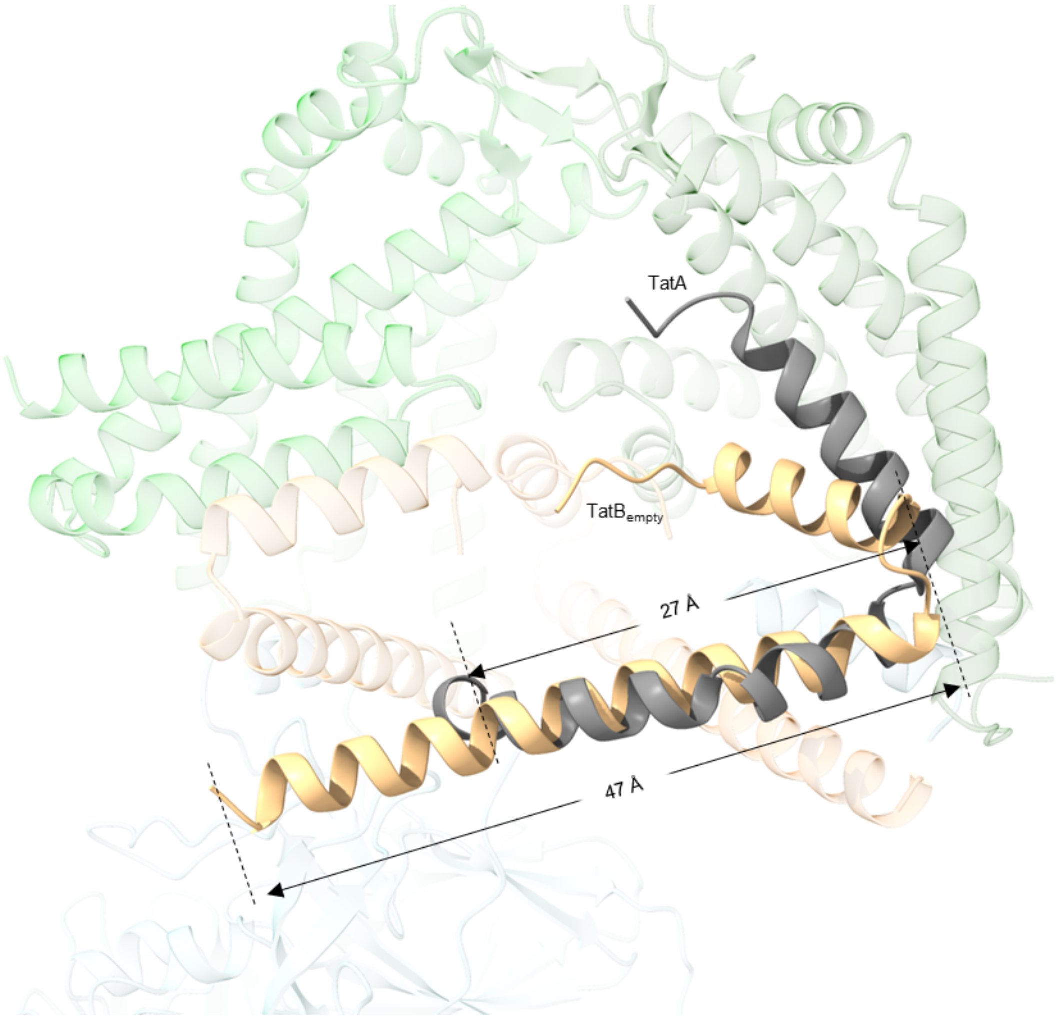


**Fig. S17. Superposition of TatA amphipathic helix with TatB$_{empty}$.**

TatA structure (PDB ID 2LZS) is shown in grey. TatB$_{empty}$ is shown in light orange. Other subunits are transparent for clarity.

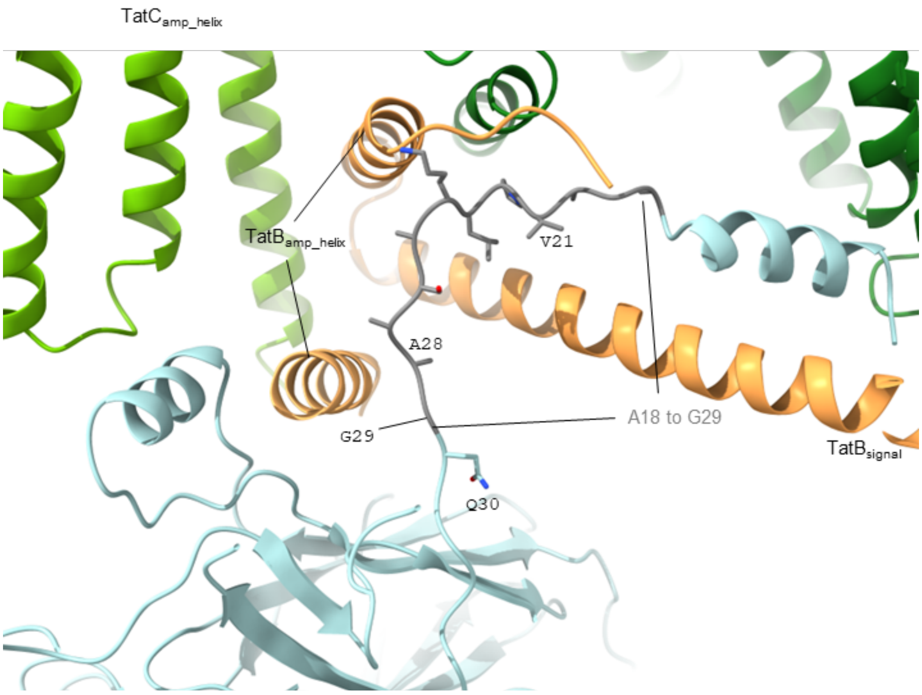


**Fig. S18. Possible modelling of missing SufI region from A18 to G29.**
Putative conformation of SufI from A18 to G29, colored in grey. The rest of the structure is our TatBC-SufI complex, colored as in Fig. 1.

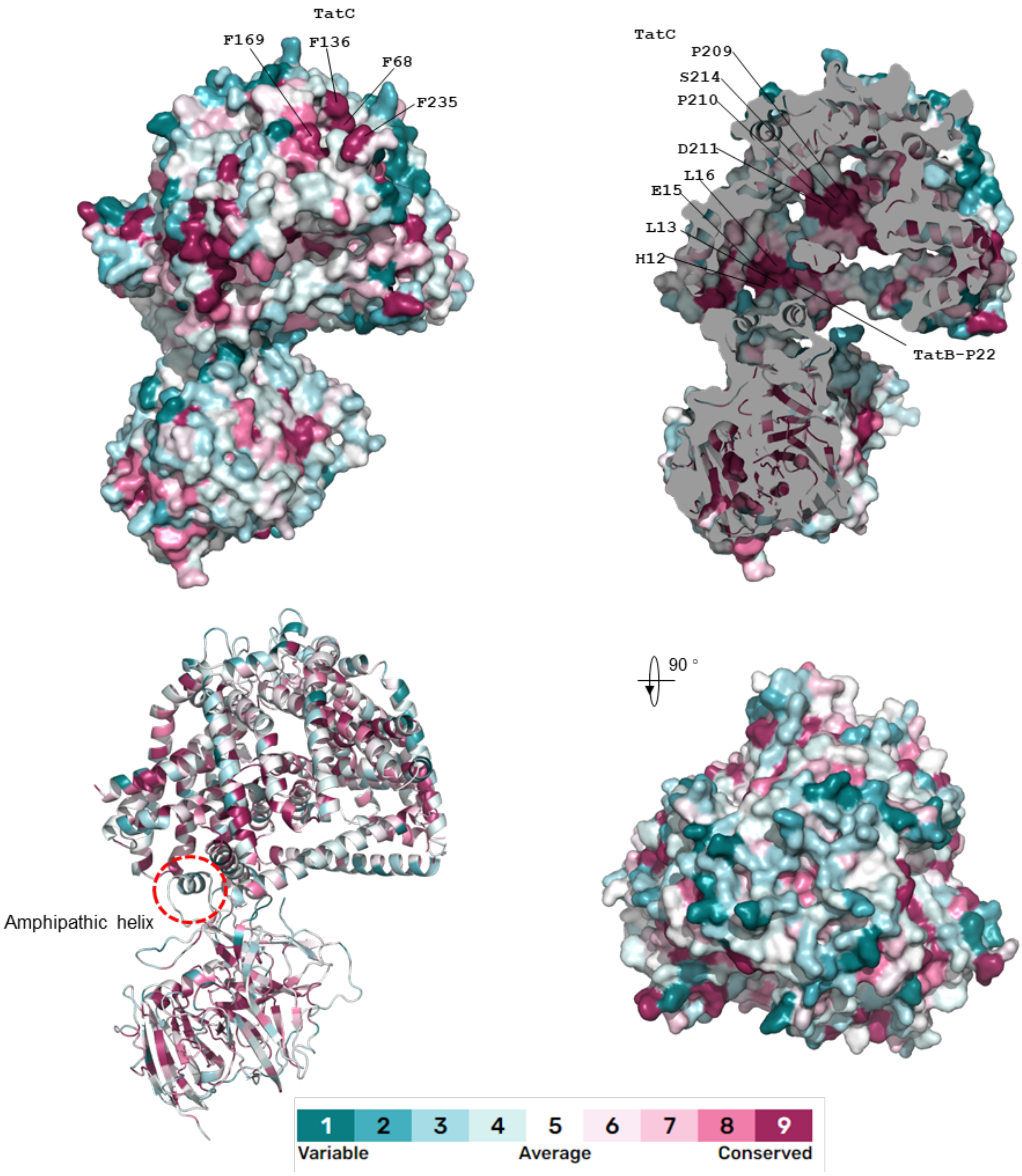


**Fig. S19. Conservation of the entire TatBC/SufI complex.**

Residues are colored according to their conservation score using ConSurf server (https://consurf.tau.ac.il/). Residues with high conservation are colored maroon and low conservation cyan. Residues with unreliable conservation scores are colored yellow, displayed on our $TatB_3C_3$-SufI structure. Inner and surface-exposed ConSurf score 9 residues are labelled. The variable short amphipathic helix SA that docks to Tat is highlighted with red dashed circle.

.

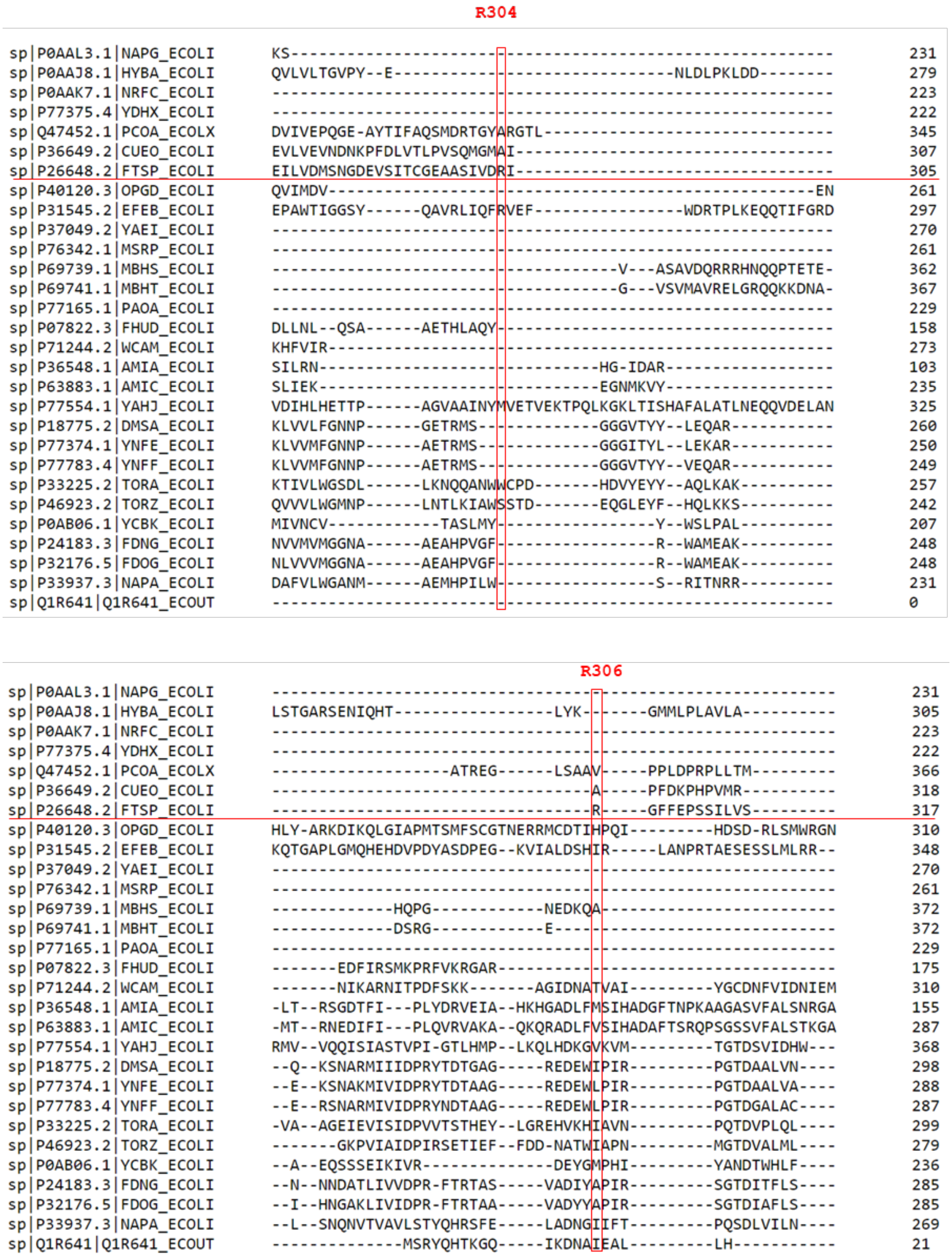


**Fig. S20. Sequence alignment of *E. coli* Tat substrates.**

The 29 known *E. coli* substrates of Tat system (56) were aligned using UniProt sequence alignment (https://www.uniprot.org/help/sequence-alignments). SufI is also known as FtsP, which is underlined by red line. R304 and R306 are highlighted.

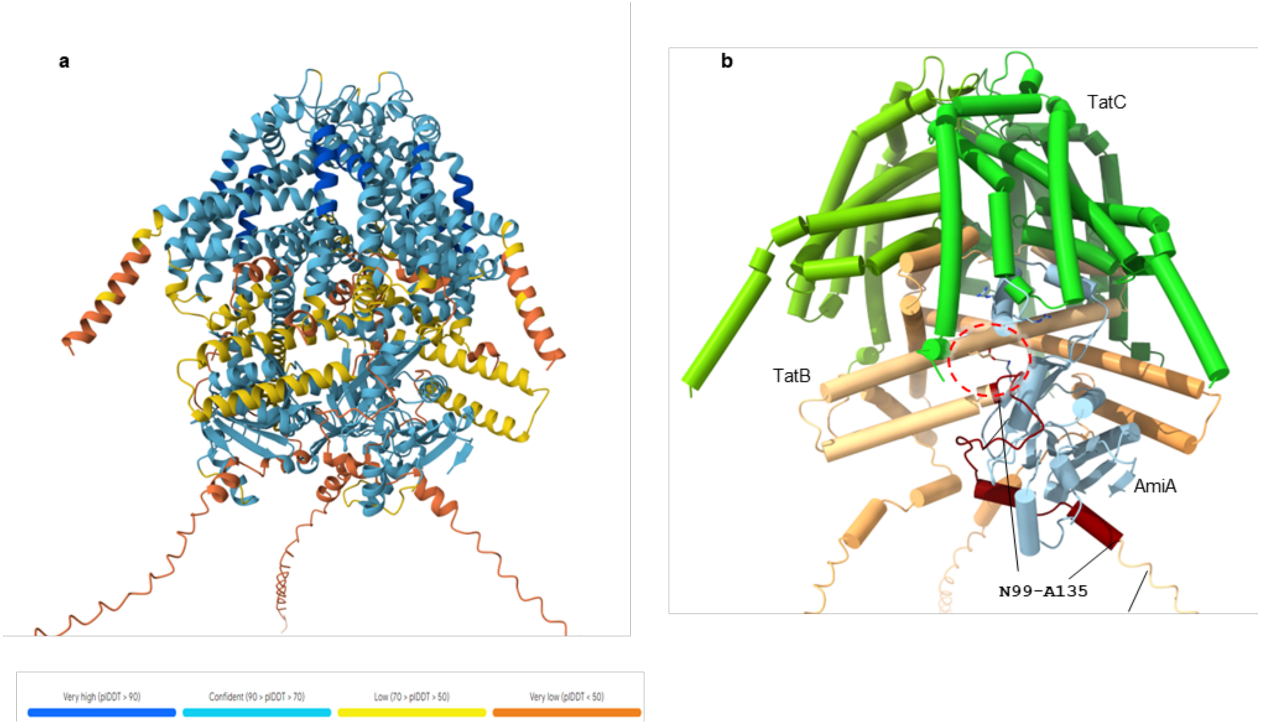


**Fig. S21. AF3-predicted model of TatBC trimer with substrate AmiA.**

Three copies each of *E. coli* TatB, TatC, and pre-AmiA were used as input. **a**, Per-atom confidence (pLDDT) of the AlphaFold prediction is color-coded from high (blue) to low (yellow to orange). **b**, Complex shown as cartoon cylinders colored by subunit. AmiA is colored cyan, and the $4^{th}$ helix of TatB N99-A135 is colored maroon. AmiA docks to TatB amphipathic helix via TatB E49 and AmiA K68 salt bridge interaction, highlighted with red dashed circle.

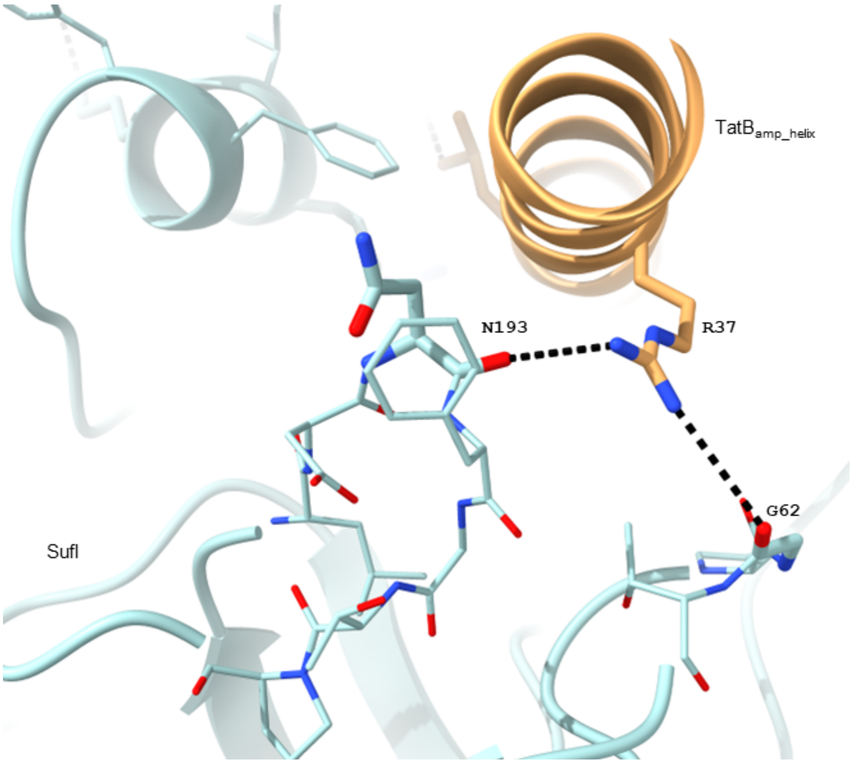


**Fig. S22. Interactions between TatB$_{amp_helix}$ and SufI.**

Backbone of SufI G62 and N193 is close to side chain of TatB R37. The black dashed lines indicate potential H-bonds with cut-off of 4 Å.

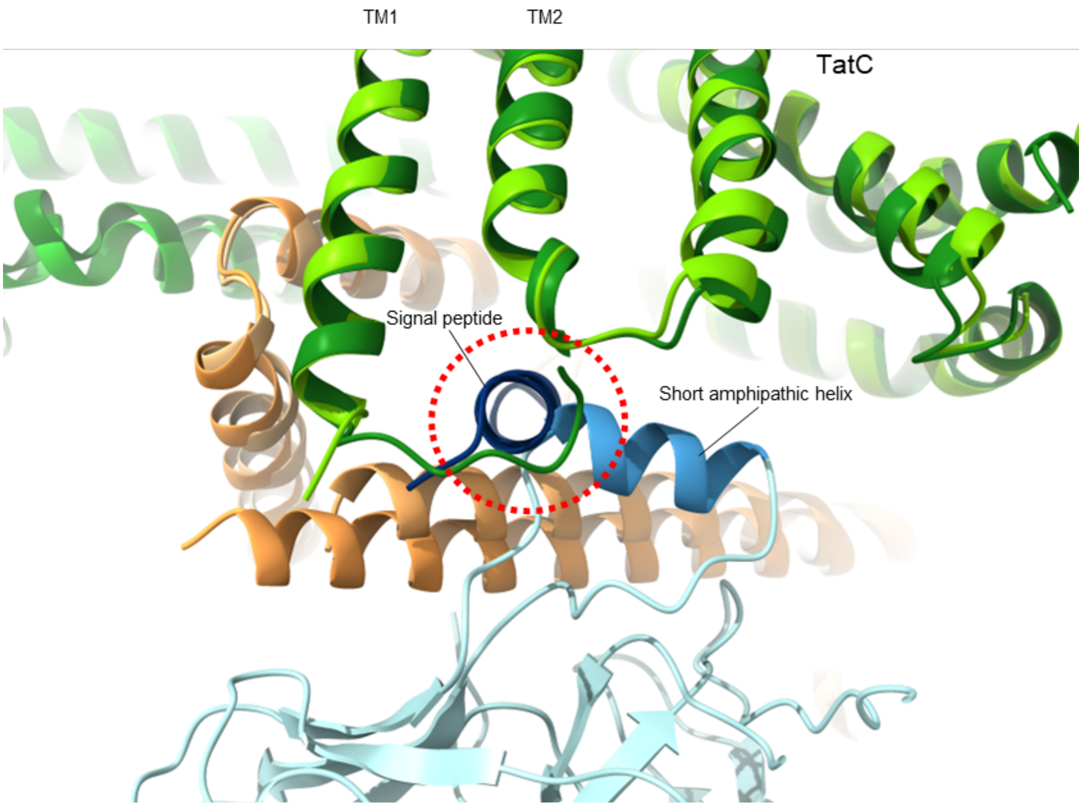


**Fig. S23. Superposition of $TatC_{amp\_helix}$ to $TatC_{signal}$.**

$TatC_{amp\_helix}$ is superposed with $TatC_{signal}$, leading to clashes between the signal peptide (dark blue) and the short helix SA of SufI (blue).

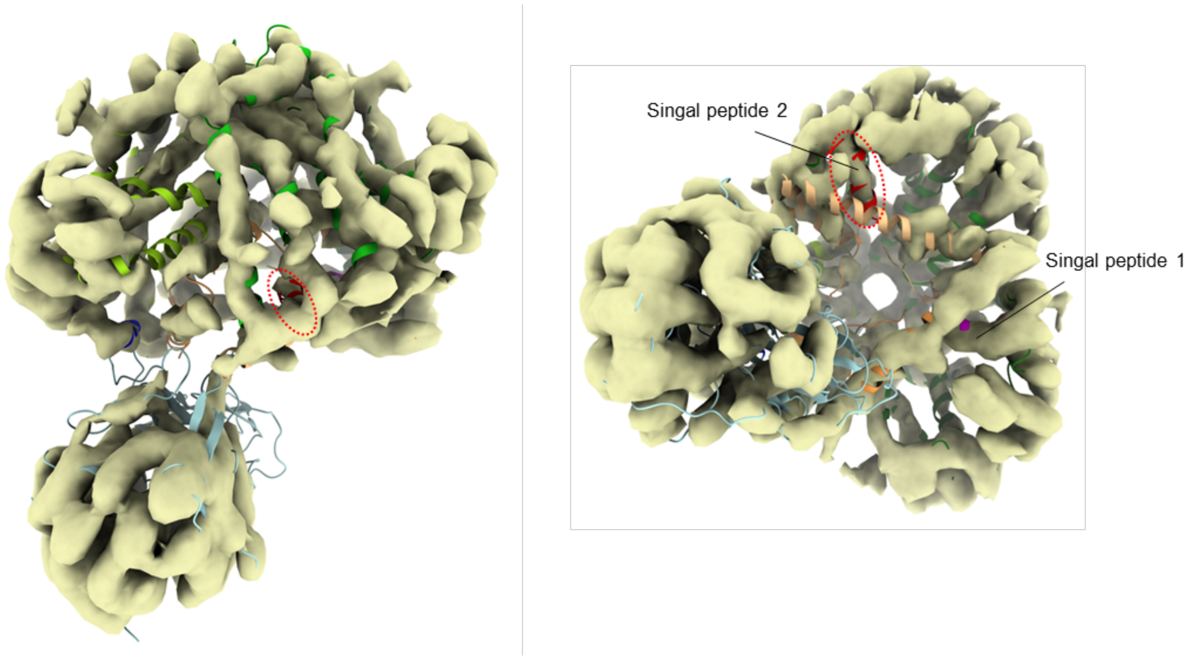


**Fig. S24. Cryo-EM density map of two SufI particles.**

A subset of 300k particles clearly containing two copies of SufI was selected and subjected to 3D refinement with mask covering Tat and just one SufI, as it provides higher resolution. Subsequent 3D classification without alignment (T=64) yielded five classes. Four out of the five classes revealed an additional density at the signal peptide binding site of $TatC_{empty}$, supporting the presence of a second bound signal peptide. This putative signal peptide is colored in red and highlighted in red dashed circle in the cytoplasmic view on the right.

| | TatBC-SufI |
|---|---|
| **Data collection and processing** | |
| Microscope | Titan Krios |
| Camera | K3 |
| Magnification | 81000X |
| Voltage (kV) | 300 |
| Electron exposure (e–/Å²) | 80 |
| Automation software | EPU |
| Number of frames | 80 |
| Defocus range (μm) | ~ -1.5 to -2.5 |
| Pixel size (Å) | 1.06 |
| Symmetry imposed | C1 |
| Number of micrographs | 10,177 |
| Initial particle images (no.) | 1.4 M |
| Final particle images (no.) | 163739 |
| Map resolution (Å) at 0.143 FSC threshold | 3.74 |
| **Refinement** | |
| Initial model used (PDB code) | Alphafold3 |
| Refinement package | Phenix_real _space_refinement |
| Model resolution (Å) at 0.5 FSC threshold | 3.9 |
| Local resolution range (Å) | 3.5-6 |
| Cross-correlation | |
| Mask | 0.79 |
| Volume | 0.78 |
| Map sharpening B factor (Å²) | -109.787 |
| Model composition | |
| Non-hydrogen atoms | 10321 |
| Protein residues | 1315 |
| B factors (Å²) (mean) | |
| Protein | 107.66 |
| R.m.s. deviations | |
| Bond lengths (Å) | 0.004 |
| Bond angles (°) | 0.793 |
| Validation | |
| MolProbity score | 1.43 |
| Clashscore | 5.15 |
| Poor rotamers (%) | 1.58 |
| C-beta deviations | 0 |
| CaBLAM outliers (%) | 0.94 |
| Ramachandran plot | |
| Favored (%) | 98.38 |
| Allowed (%) | 1.54 |
| Disallowed (%) | 0.08 |

**Table S1. Data processing and model refinement statistics.**

| Subunit name | Chain | Range built/total residues | Un-modelled residues | % modelled residues |
|---|---|---|---|---|
| TatC | A | 2-237/258 | 1, 238-258 | 91.5 |
| TatC | B | 7-235/258 | 1-6, 236-258 | 89.1 |
| TatC | C | 8-235/258 | 1-7, 236-258 | 88.4 |
| TatB | D | 1-58/171 | 59-171 | 33.9 |
| TatB | E | 1-52/171 | 53-171 | 30.4 |
| TatB | F | 2-57/171 | 1, 58-171 | 32.7 |
| Sufi | H | 3-17, 30-470/470 | 1-2, 18-29 | 97 |
| $TatB_3C_3$-SufI | | 1315/1757 | | 74.8 |

**Table S2. Modelling of Tat complex.**

| Substrate | Docking interactions observed (Y/N) | Singal peptide bound (Y/N) | TatB C-terminus wraps substrate (Y/N) |
|---|---|---|---|
| P36548_AMIA | Y | Y | Y |
| P0AB06_YCBK | Y | Y | Y |
| P07822_FHUD | N | Y | Y |
| P40120_OPGD | N | Y | Y |
| P18775_DMSA | Y | Y | Y |
| P33225_TORA | N | Y | Y |
| P77165_PAOA | Y | Y | N |
| P0AAL3_NAPG | Y | N | Y |
| P69739_MBHS | N | N | Y |
| P0AAJ8_HYBA | N | N | Y |

**Table S3. AF3 models of $TatB_3C_3$ with bound substrates.**

The substrate sequences were obtained from UniProt (https://www.uniprot.org/), with the corresponding accession IDs listed before the substrate names. Three copies of substrates, along with three copies each of *E. coli* TatB and TatC were used as input for AF3.

| Sites | Interactions in the $TatB_3C_3$-SufI complex structure | C. score | Functional effect of mutations | Source |
|---|---|---|---|---|
| **TatC** | | | | |
| H12 | H-bond with SufI-R5, π–π Stacking interaction with SufI-F8 | 9 | H12A reduced transportation efficiency | (Buchanan et al., 2002) |
| E15 | Salt bridge with SufI-R5 | 9 | E15A reduced transportation efficiency | (Buchanan et al., 2002) |
| R17 | H-bond to TatB-V18 backbone oxygen. Neutralizes negative dipole of TatB TM1 C-terminus | 9 | R17A reduces transport activity; R17K reduces activity less than R17A; R17A leads to reduced complex size on BN-PAGE | (Allen et al., 2002) (Buchanan et al., 2002) (Behrendt & Brüser, 2014) |
| R19 | H-bond to TatC-Q90 | 9 | R19A, R19K reduce somewhat transportation activity | (Buchanan et al., 2002) |
| I60 | Sealing complex at the periplasmic side | 8 | I60N loses transportation activity, Introducing an alanine after I60 or deletion of A61 also abolishes transportation | (Kneuper et al., 2012) |
| T62 | Sealing complex at the periplasmic side | 8 | Deletion of T62 abolishes transportation | (Kneuper et al., 2012) |
| D63 | H-bond to adjacent TatC-S153, provides charge for inner cavity at periplasmic side | 5 | D63V abolishes translocation | (Kneuper et al., 2012) |
| V64 | TatC-TatC interface; Hydrophobic interactions with adjacent TatC F134, L149, Y154, F157 | 7 | V64E abolishes translocation | (Kneuper et al., 2012) |
| F68 | TatC-TatC interface; Hydrophobic interactions with adjacent TatC F136, L137 | 9 | F68S abolishes transportation | (Kneuper et al., 2012) |
| K73 | Salt bridge with adjacent TatC-D211, pH sensor? | 8 | Deletion of K73 abolishes transportation; reduced amount of Tat | (Kneuper et al., 2012) |
| P85 | In-helix proline, bending TM2 | 9 | P85A reduces the translocation effiency | (Buchanan et al., 2002) |
| Q90 | H-bond with R19 | 9 | Q90A reduces translocation | (Buchanan et al., 2002) |
| W92 | Hydrophobic interactions with TatC TM3 residues at cytoplasmic side | 8 | W92A reduces translocation efficiency somewhat | (Buchanan et al., 2002) |
| F94 | Hydrophobic interactions with SufI-F8, I9; cation-π interactions with R5 | 9 | Mutations to A, L, S abolish Tat function; mutations to Y no effect on Tat function | (Buchanan et al., 2002) (Kneuper et al., 2012) |
| P97 | Linker between TM2 and TM3 | 9 | P97A reduces transportation efficiency | (Buchanan et al., 2002) |
| L99 | Hydrophobic interactions with SufI-V302 | 9 | L99A reduces transportation efficiency | (Buchanan et al., 2002) |
| Y100 | Cation-π interactions with SufI-R306 | 8 | Y100A reduces transportation efficiency | (Buchanan et al., 2002) |
| E103 | Salt bridge with SufI-R6 and SufI-R306 | 9 | Mutations to A, D, Q, R abolish transportation; to G, K abolish growth on SDS | (Buchanan et al., 2002) (Kneuper et al., 2012) |
| R105 | No obvious interactions | 7 | R103A reduces translocation efficiency a little | (Buchanan et al., 2002) |
| Y126 | Hydrophobic interactions with surrounding hydrophobic residues, and H-bond with L44, stabilizes the TatC monomer | 8 | Y126A reduces transportation efficiency | (Buchanan et al., 2002) |
| L137 | Hydrophobic interactions with adjacent F68 and loop | 7 | L137H abolish Tat translocation | (Kneuper et al., 2012) |
| D150 | H-bond to TatC-T62 (sealing residue), provides charge for inner cavity at periplasmic side | 7 | D150G, D150V abolish transportation activity | (Kneuper et al., 2012) |
| Y154 | Hydrophobic interactions with adjacent TatC-V64 | 9 | Y154H has no obvious effect on Tat function, Y154A no obvious effect on Tat complex size | (Kneuper et al., 2012) (Behrendt & Brüser, 2014) |
| F157 | Hydrophobic interactions with adjacent TatC-V64 | 8 | F157Y has no obvious effect on Tat function | (Kneuper et al., 2012) |
| E170 | No obvious interactions, but provides charge for inner cavity | 8 | E170A does not affect signal binding, but reduces transport efficiency | (Holzafer et al, 2007) (Buchanan et al., 2002) |
| P172 | Proline in TM4 | 9 | P172A reduce translocation efficiency a little | (Buchanan et al., 2002) |

| | | | | |
|---|---|---|---|---|
| M205 | H-bond with TatB-E8 | 9 | M205A leads to reduced transportation; M205R more retarded transportation; both M205RA still bind TatB and TatA | (Alcock et al., 2016) |
| T208 | H-bond with TatB-E8 | 9 | T208A reduces translocation efficiency; TatB and TatA still bind | (Alcock et al., 2016) |
| D211 | Salt bridge with adjacent TatC-K73 from TM5, pH sensor? provides charge for inner cavity | 9 | Mutation to A abolishes Tat function, to E and N results in reduced efficiency | (Buchanan et al., 2002) |
| S214 | H-bond with D211 (TM5,6 linker residue) | 9 | S214A reduces translocation efficiency a little | (Buchanan et al., 2002) |
| Q215 | H-bond with TatB-E8 | 9 | Q215A reduces transportation efficiency, TatB and TatA still bind | (Alcock et al., 2016) |
| E227 | Salt bridge with TatC-R193 | 9 | E227A reduces translocation efficiency a bit | (Buchanan et al., 2002) |
| **TatB** | | | | |
| F2 | Hydrophobic interactions with adjacent TatB-F6 | 9 | TatB F2L with F6L restore transportation in TatC M205C; shows TatB dimers when crosslinked | (Blümmel et al., 2015) (Kneuper et al., 2012) |
| D3 | H-bond with TatC-T208 backbone oxygen | 9 | D3A mutation together with E49A has weaker effect on TatB/TatC binding than E8A | (Fröbel et al., 2019) |
| I4 | Possible hydrophobic interaction with TatC-L206 | 9 | shows TatB multimers when crosslinked | (Blümmel et al., 2015) |
| F6 | Hydrophobic interactions with adjacent TatB-F2 | 9 | shows TatB multimers when crosslinked | (Kneuper et al., 2012) (Blümmel et al., 2015) |
| E8 | H-bonds with TatC-MTQ polar cluster | 9 | E8A abolishes transportation, reduces TatB/TatC binding | (Alcock et al., 2016) |
| F13 | $TatB_{signal}$-F13 form hydrophobic interactions with adjacent TatC TM1, while $TatB_{amp\_helix}$ doesn't have this interaction | 6 | F13Y show un-specific transportation without signal peptide | (Huang et al., 2017) |
| E49 | Salt bridge with SufI-R304; close to TatC N-terminus | 9 | Single E49A mutation does not affect the crosslinking between substrate signal peptide and TatB, but E49A together with E53A diminish such crosslinking. | (Fröbel et al., 2019) |

**Table S4. Structural interpretation of the functional effects of mutations introduced into E. coli TatC and TatB proteins.**

"Adjacent" indicates that interacting residue comes from the neighbouring subunit. C. score indicates conservation score calculated with ConSurf server, from 1 to 9 (variable to conserved).
